# scSVAS: CNV clonal visualization online platform for large scale single-cell genomics

**DOI:** 10.1101/2021.06.10.437122

**Authors:** Lingxi Chen, Yuhao Qing, Ruikang Li, Chaohui Li, Hechen Li, Xikang Feng, Shuai Cheng Li

## Abstract

The recent advance of single-cell copy number variation analysis plays an essential role in addressing intra-tumor heterogeneity, identifying tumor subgroups, and restoring tumor evolving trajectories at single-cell scale. Pleasant visualization of copy number analysis results boosts productive scientific exploration, validation, and sharing. Several single-cell analysis figures have the effectiveness of visualizations for understanding single-cell genomics in published articles and software packages. However, they almost lack real-time interaction, and it is hard to reproduce them. Moreover, existing tools are time-consuming and memory-intensive when they reach large-scale single-cell throughputs. We present an online visualization platform, scSVAS, for real-time interactive single-cell genomics data visualization. scSVAS is specifically designed for large-scale single-cell analysis. Compared with other tools, scSVAS manifests the most comprehensive functionalities. After uploading the specified input files, scSVAS deploys the online interactive visualization automatically. Users may make scientific discoveries, share interactive visualization, and download high-quality publication-ready figures. scSVAS provides versatile utilities for managing, investigating, sharing, and publishing single-cell copy number variation profiles. We envision this online platform will expedite the biological understanding of cancer clonal evolution in single-cell resolution. All visualizations are publicly hosted at https://sc.deepomics.org.

## Introduction

With the explosion of single-cell DNA sequencing (scDNA-Seq) in cancer analysis, it becomes increasingly crucial to accurately calling the somatic variants in single-cell resolution. Compared with the traditional bulk DNA-Seq method, the scDNA-Seq exhibits several advantages. Although the former studies have contributed perspicacity into tumor biology, it is restricted to offering the mixed signals of tumor cells or clones which hold genotype diversity, leading to the mask of intra-tumor heterogeneity (1). The coming of scDNA-Seq addresses this concern soundly (2). Researchers can conquer the obstacles of bulk profiling to mark intra-tumor heterogeneity (3), distinguish tumor subclones (1), build clonal lineage and metastasis (4), and infer the cause of therapeutic resistance (5). Analyses for scDNA-Seq may go into three categories: (i) applying dimension reduction techniques to decipher the clonal substructure; (ii) building the clonal lineage within a tumor (2), among multi lesions (4), or along time (6); (iii) resolving mutation co-occurrence and mutual exclusivity across subclones (7).

Pleasant visualization of single-cell analysis results boosts productive scientific exploration, validation, and sharing (8). Several single-cell analysis figures have the effectiveness of visualizations for understanding single-cell genomics been demonstrated in published articles and software packages. Nevertheless, those pictures almost lack real-time interaction, and it is hard for researchers to prepare codes to reproduce them. Moreover, with the stride of high throughput scDNA sequencing, the scale of sequenced cells escalates exponentially, aka, hundreds or thousands of cells at a time (3, 9, 10). Efficient visualization of single-cells with a large (e.g. 1k × 5k) size is critical for scientific interpretation. Plotting using R or Python packages, or existing single-cell visualization tools are incredibly time-consuming and memory-intensive when it reaches thousands of cells and thousands of genomic regions.

Therefore, we present an online platform scSVAS (https://sc.deepomics.org) for aesthetically-pleasing, real-time interactive, and user-friendly single-cell copy number variation (CNV) visualization (Fig. 1A, Supplementary Fig. S1-S11). To our knowledge, scSVAS is the first online platform specialized for large-scale scDNA CNV visualization. We offer the users an editor to upload the required upstream analysis output and customize the display settings (Supplementary Fig. S12, Supplementary Manual). We provide an interactive tooltip to display vital information for each visualization object, assisting users in making scientific discoveries effectively. scSVAS is code-free for users. All visualizations are downloadable in high-quality publication-ready format. We support dark and light theme for visualization and offers a collection of scientific (SCI) journal color palettes. In all, compared with existed tools Ginkgo (11), E-Scape (12), and 10x Loupe scDNA Browser (3), scSVAS manifest the most comprehensive functionalities (Fig. 1B-C, Table 1).

**Table 1.** Key functionalities of scSVAS visualizations. ✓ is marked if benchmark tools Ginkgo (11), E-Scape (12), or 10x Loupe (3) supports the described functionality, -otherwise.

| scSVAS Applications | Key Functionalities | Ginkgo (11) | E-Scape (12) | Loupe (3) |
| --- | --- | --- | --- | --- |
| Common features | Online platform | ✓ | - | - |
|  | Interactive plot | - | ✓ | ✓ |
|  | Code free | ✓ | - | ✓ |
|  | SCI color palette | - | - | - |
|  | Dark-light theme | - | - | - |
|  | Downloadable high-quality figures | ✓ | ✓ | ✓ |
| CNV View | Cell CNV heatmap | ✓ | ✓ | ✓ |
|  | Cell meta heatmap | - | - | ✓ |
|  | Zoomable cell dendrogram | - | - | ✓ |
|  | Group CN heatmap/stairstep | - | - | - |
| CNV Heatmap | Cell CNV heatmap | ✓ | ✓ | ✓ |
|  | Cell meta heatmap | - | - | ✓ |
|  | Zoomable cell dendrogram | - | - | ✓ |
|  | Gene Track | - | - | ✓ |
|  | RepeatMasker Track | - | - | - |
|  | Zoomable genomic region | - | - | ✓ |
|  | Search local genomic region | - | - | ✓ |
| Cell Phylogeny | Zoomable (topdown/circular) cell dendrogram | - | - | - |
|  | Cell-gene CNV heatmap | - | - | - |
|  | Cell-bin CNV heatmap | - | - | - |
| Ploidy Stairstep | Cell/group/total ploidy stairstep | - | - | - |
| Ploidy Distribution | Cell/group/total ploidy distribution | - | - | - |
| Embedding Map | 2D embedding plot | - | - | - |
|  | Scatter plot/hexagonal binning | - | - | - |
|  | Color annotation: cell density/gene CNV/meta information | - | - | - |
| Time Lineage | Evolution fishplot with different layout | - | ✓ | - |
|  | Fishplot with bullet shape | - | - | - |
|  | Circular/acute lineage tree | - | - | - |
|  | Cell ensemble presentation | - | - | - |
| Space Lineage | Subclone lineage tree | - | ✓ | - |
|  | Lesion lineage tree | - | - | - |
|  | Zoomable anatomy image | - | - | - |
|  | Movable lesion pointer | - | - | - |
| Space Prevalence | Subclone lineage tree | - | ✓ | - |
|  | Lesion lineage tree | - | - | - |
|  | Clonal frequency matrix/bipartite graph between subclone and lesion | - | - | - |
| Clonal Lineage | Subclone lineage tree | - | - | - |
|  | Group CNV heatmap | - | - | - |
|  | Cell ensemble presentation | - | - | - |
|  | Clonal stairstep comparison | - | - | - |
|  | Customized gene selection and gene set annotation | - | - | - |
| Recurrent Event | Recurrent stairstep comparison | - | - | - |
|  | Recurrent focal gain/loss | - | - | - |
|  | Customized gene selection and gene set annotation | - | - | - |

**Fig. 1.**
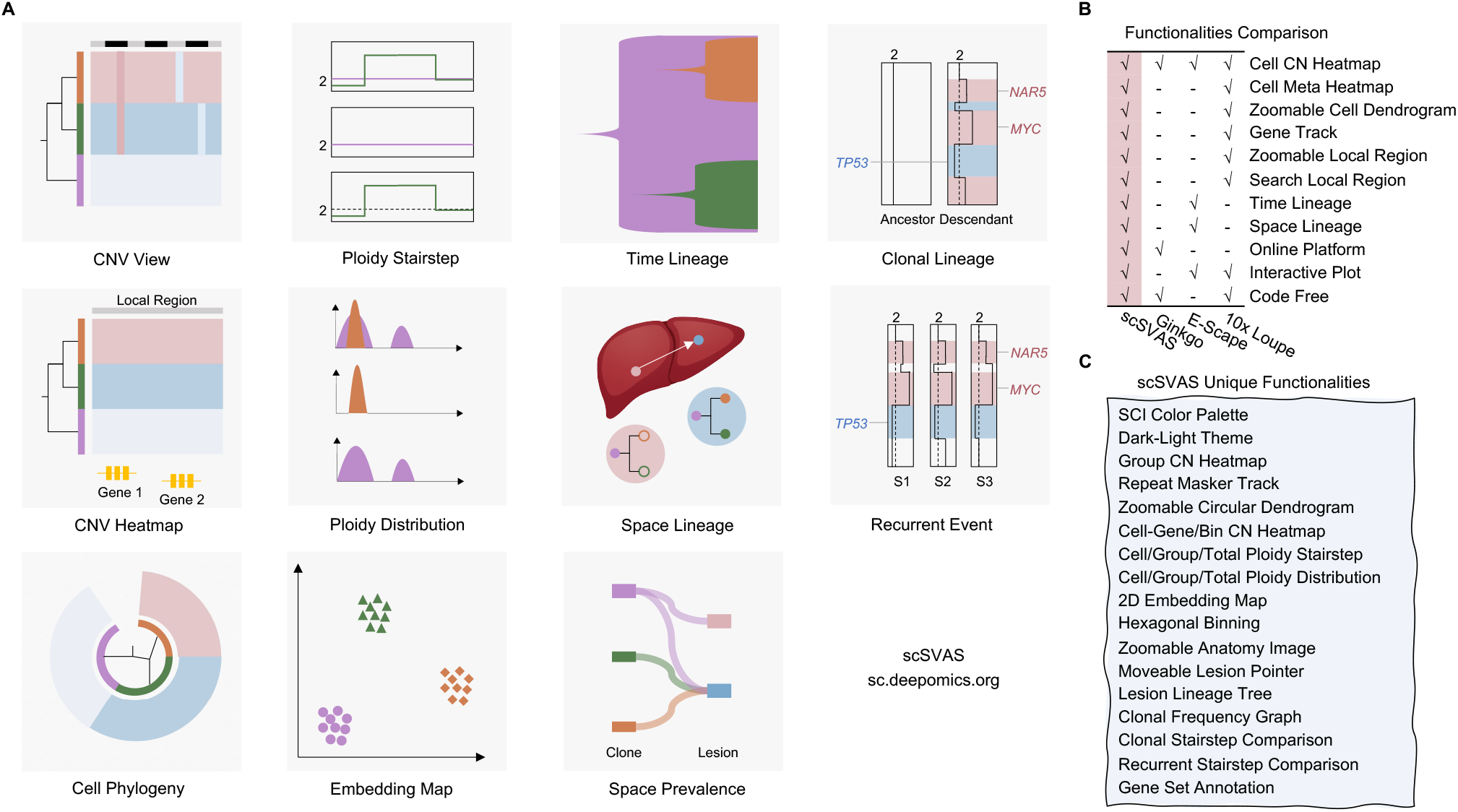
A Thumbnails of 11 scSVAS visualization applications. **B** Functionalities comparison between scSVAS, Ginkgo (11), E-Scape (12), and 10x Loupe scDNA Browser (3). **C** scSVAS unique features compared with other two tools. CN: Copy Number. SCI: Scientific Journal. 2D: 2 Dimension. Freepik and macrovector / Freepik design the cartoon anatomy image, we acknowledge for the free license.

## Results

### Visualization of single-cell CNV landscape and phylogeny

CNV Heatmap is an intrinsic way to visualize the landscape of single-cell CNV profiles in several pieces of literature (7, 13, 14). Cooperating scSVAS offline pipeline, we build a web interface CNV View as demonstrated in Supplementary Fig. S1 and Table 1. CNV View exhibits the copy number of single-cells across the entire genome with single-cells as rows and genomic regions as columns (Supplementary Fig. S1A). If users offer the cut dendrogram file, a zoomable cut dendrogram will be displayed on the left (Supplementary Fig. S1A,B,C). The cell meta-annotation heatmap will be displayed on the left if users provide single-cell meta information (Supplementary Fig. S1A,D). In addition, the aggregate subgroup CNV heatmap and stairstep will also be listed in the bottom layers (Supplementary Fig. S1A,E,F).

CNV Heatmap (Supplementary Fig. S2) extends CNV View by supporting extra genome region zooming, local region search, and local region annotation (repeats and genes). There is a genome zoom slider located at the bottom of CNV heatmap (Supplementary Fig. S2A). Users can drag the slider to zoom in and out on the genome region. Users can also search for a local genome region. If the local genome region is less than 5M base pairs, an annotation layer including “Repeat track” and “Gene track” will be displayed (Supplementary Fig. S2B).

Besides, we build Cell Phylogeny (Supplementary Fig. S3) to concentrates on the zoomable cut dendrogram of single-cells. It offers two layouts (top-down and circular), and supports cell-gene or cell-region CNV as meta-annotation.

### Visualization of single-cell ploidy profile

The tumor CNV in different regions may encounter dramatic gain or loss (15). The ploidy line plot along the chromosomes can visually show the heterogeneity between tumor subclones by combining genomic coordinates. By collapsing the single-cells in the same tumor subclones into one observation, we can infer the pseudo-bulk ploidy of each subclone. Since cancer CNV ploidy line alters along chromosomes, we call it the “stairstep plot”. Besides, visualization of ploidy distribution reveals the intra-tumor heterogeneity as well. Thus, we developed two web interface Ploidy Stairstep and Ploidy Distribution demonstrated in Supplementary Fig. S4 and Supplementary Fig. S5, respectively. The layout is a matrix of the ploidy stairstep or density plot. The column lists categorical meta-labels available in uploaded file by default. The first row exhibits the ploidy of all subgroups for specific categorical meta labels in an aggregate form. The second line displays the collapsed ploidy of total single-cells as the way for bulk sequencing. Then, the subsequent rows list the ploidy of all available subclones, respectively.

### Visualization of single-cell 2D-embedding

High-dimensional data could be challenging to visualize. Reducing data into two dimensions is essential for representing the inherent structure of the data. In terms of large-scale scDNA data, conventional 2D scatter plots may disguise essential information. Embedding Map (Supplementary Fig. S6) defeats this overplotting obstacle utilizing hexagonal binning (16, 17), which also has benefits on time and memory complexity. We offer different strategies to color the single-cell data point. If the “hexagon mode” is activated, the embedded cells colored with density will be displayed. With gene CNV profiles, the embedded cells can be annotated by the specified gene’s copy number. Moreover, all categorical meta labels available will be used as color schemes by default. We also provided the scSVAS offline pipeline which offers diverse embedding techniques (e.g., PCA, UMAP (18), TSNE, PHATE (19), and DeepMF (20)) to generate the input files.

### Visualization of clonal dynamics in time and space

Single-cell genomics data help researchers to resolve the clonal lineage within a tumor (2), among multi lesions (4), or along time (6). Here, we build web interfaces Time Lineage, Space Lineage, and Space Prevalence to visualize the evolution dynamics between tumor subclones over time and space.

Time Lineage (Supplementary Fig. S7A) is composed of fishplot, lineage tree, and cellular ensemble. The fishplot conceptually manifests the proportion of tumor subclones at different tumorigenesis stages along time. We use the bezier curve to fit the trend of subclones over time with two distinct head shapes (bullet and onion) (Supplementary Fig. S7B). The lineage tree exhibits the evolutionary relationship between tumor subclones, three different shapes (topdown normal, circular normal, and circular acute) are served (Supplementary Fig. S7C). The cellular ensemble is an abstract aesthetic presenting the tumor’s cellular prevalence at a certain time point (Supplementary Fig. S7D).

Space Lineage (Supplementary Fig. S8A) visualizes the clonal dynamics across multiple lesions. We offer subclone and lesion lineage trees. We also provide a zoomable human anatomy atlas (license-free) (Supplementary Fig. S8B-C). Users can select different anatomy images from cartoon anatomy atlas (male, female, brain, intestines, kidney, liver, lung, pancreas, stomach, thymus, thyroid, urinary system, male reproductive system, and female reproductive system). Instead of the arduous lesion position precalculation required in E-Scape (12), we provide an image pointer for each lesion. Users can right-click to active it and drag the pointer to the exact lesion position (Supplementary Fig. S8D).

Space Prevalence (Supplementary Fig. S9) provides the sub-clone and lesion lineage trees and visualizes the clonal prevalence across subclones and lesions utilizing matrices and bi-partite graphs.

### Visualization of CNV shift across subclone

To further illustrate how CNVs change through the clonal evolution history, we built the web interface Clonal Lineage (Supplementary Fig. S10A), which is composed of lineage tree, sub-clone CNV heatmap, cellular ensemble, lineage tree branch, parent/child CNV stairstep, and gene box. The lineage tree shares four different types: topdown normal, circular normal, circular acute, and fishplot as demonstrated in Supplementary Fig. S10B. The group CNV heatmap displays the averaged copy number profiles of subclones. The lineage tree branch displays the number of gain and loss regions for each tree branch. The stairsteps depict the detailed CNV shift from the parent node to the child node annotated with deleted and amplified genes. We offer MsigDB (21) and customized gene set annotation with stacked and donut modes (Supplementary Fig. S10C). Users can click the gene to direct to the GeneCards (22) website page. One can click the lineage tree branch to check the different CNV shifts and study the clonal dynamics from multiple timepoint visualization of clonal lineage across time.

### Visualization of focal gain and loss across cohort

Recurrent focal CNV across a batch of tumor samples may be the possible driver CNV (23). Here, Recurrent Event (Supplementary Fig. S11) provides the interactive and real-time visualization of focal gains and losses across multiple sub-clones and samples. In the middle, it displays the CNV stairsteps of all subclones across samples. The left and right gene boxes show the recurrent gain and loss genes, respectively. Similar to Clonal Lineage, customized gene selection, and gene set (self-customized and MsigDB) annotation is provided. Besides, the bottom layers demonstrate available meta-sample information.

## Discussion

Visualization of single-cell CNV data plays an essential role in sharing scientific results and acting as an auxiliary tool for data investigation. Herein, we built an online visualization platform, scSVAS, that offers user-friendly and real-time interactive single-cell genomics data visualization. scSVAS is specifically designed for large-scale single-cell analysis. scSVAS is code-free; users have loads of choices to customize the visualization with simple mouse operations and export the visualizations into figures. We have deployed eleven web visualization interfaces, including CNV View, CNV Heatmap, Cell Phylogeny, Ploidy Stairstep, Ploidy Distribution, Embedding Map, Time Lineage, Space Lineage, Space Prevalence, Clonal Lineage, and Recurrent Event. In addition, we provided informative user manual and demo cases.

Existing tools visualize the single-cell genomics and copy number evolution include Ginkgo (11), E-Scape (12), and 10x Loupe scDNA Browser (3) (Table 1). Ginkgo is an online single-cell CNV caller which only provides static cell CNV heatmap figures and is incompatible with large-scale single cell data. E-Scape provides interactive visualization of cell CNV heatmap and clonal evolution (in fishplot and sub-clone lineage tree) across time and space. As it is powered with R Shiny, it requires R programming skills, and there exits memory and time deficiency when dealing with thousands of single cells. 10x Loupe scDNA Browser focuses on interactively visualizing the CNV profile of single-cell genomics, including cell CNV heatmap, cell meta heatmap, zoomable cell dendrogram, gene track, zoomable genomic region, and search local genomic region. However, it only accepts the data generated from the 10x genomics CNV solution and pipeline. With the aid of Oviz (8) visualization framework, scSVAS platform not only supports all functionalities which competitors own but also allows more exploratory functions by using informative tooltips, simultaneous highlighting, zoom-in/out, mouse drag-in/drag-out, etc. (Fig. 1B-C and Table 1). In particular, both 10x Loupe scDNA Browser and scSVAS (CNV View and CNV Heatmap) show cell CNV heatmap with cell metadata annotation, scSVAS offers users options to sort the cells according to meta-label. Moreover, scSVAS provides RepeatMasker annotation when display local genomic regions. In another case, both E-Scape and scSVAS (Space Lineage) display clonal evolution dynamics across space, with each lesion lineage tree pointing to the exact position of the anatomy image. Unfortunately, E-Scape demands users input the coordinates of lesion positions, while the accurate lesion coordinates are tedious for users to measure. scSVAS releases users from the tiresome estimation with simple mouse operations, allowing users to activate the lesion pointer and move it to the ideal position on the anatomy image. Furthermore, we provide a zoomable human anatomy atlas (license-free) and extra zoom-in/out functionalities. In addition, scSVAS contributes a more comprehensive set of single-cell CNV analysis results, including top-down and circular cell phylogeny, CNV ploidy stairstep and distribution, CNV embedding map, CNV shift along clonal lineage, subclone-level recurrent CNV, thus affording a one-stop service for single-cell CNV clonal visualization.

We intend to continuously improve and maintain scSVAS to ensure that the platform is user-friendly and cutting-edge. We will add more annotation databases such as dbSNP and TCGA focal CNV. Currently, scSVAS are focusing on visualizing the intra-tumor heterogeneity and evolution history based on CNV. We will extend it to single nucleotide variation (SNV) and structure variation (SV) in the close future. Collectively, scSVAS provides versatile utilities for managing, investigating, sharing, and publishing single-cell copy number variation profiles. We envision this online platform will expedite the biological understanding of cancer clonal evolution in single-cell resolution.

## Acknowledgement

We thank Miss Xinye Zhang for her assistance in drawing the application thumbnail illustration. Freepik and macrovector / Freepik design the anatomy images used in Fig. 1 and Space Lineage interface, we acknowledge for the free license.

## Contributions

S. C. L. supervised the project. L. C., Y. Q., R. L., and C. L. implemented the visualization applications. H. L. provided the back-end and front-end framework of the server. L. C. and X. F. prepared the demo data. L. C. wrote the manuscript. S. C. L revised the manuscript. All authors read and approved the final manuscript.

## Data availability

All data used for visualization is available at https://docsc.deepomics.org.

## Software availability

The offline pipeline are available at https://github.com/paprikachan/scSVAS.

## Supplementary Note 1: Supplementary Methods

### 1.1. Online platform framework

scSVAS is deployed on a remote CentOS 7.4 server with 128 GB memory and 60 TB storage. We utilized Ruby on Rails (v5.2.3), Apache (v2.4.6), and PostgreSQL (v12.3) as backend framework; HTML5, Vue.js (v2.6.10), and Oviz (1) (https://oviz.org), an in-house visualization framework written in TypeScript, as frontend support. Supplementary Table S1 listed the full technology stack of scSVAS.

### 1.2. Offline pipeline

For each visualization application, users need to prepare the input files (Supplementary Fig. S12). The input format is specified in https://docsc.deepomics.org.

Most of the customized file formats are easy to generate from upstream CNV analysis. At this moment, equipped with offline pipeline, scSVAS supports output files from scDNA CNV calling tools 10X cellranger-cnv (2), SCYN (3), and SeCNV. Users can use the provided scripts to generate the required files locally and then upload them to the corresponding visualization pages. The offline pipeline are available at https://github.com/paprikachan/scSVAS.

### 1.3. Demo data

All online analyses provide demo files for an instant preview, which available at Editor on each application page. Supplementary Table S2 demonstrates the demo datasets currently adopted. For CNV View, CNV Heatmap, Cell Phylogeny, Ploidy Stairstep, Ploidy Distribution, Embedding Map, and Clonal Lineage, the raw data of TNBC_T10 and TNBC_T16 were obtained from Navin *et al*. (4) and processed by SCYN (3) and scSVAS offline pipeline. The raw data of 10x_COLO829 was obtained from Velazquez *et al*. (5) and processed by scSVAS offline pipeline. For Time Lineage, we obtained the input files of AML (6) from Smith *et al*. (7), and HGSOC_P7 (8) from Smith *et al*. (7) For Space Lineage and Space Prevalence, we obtained HGSOC_P1 and HGSOC_P7 (8) from Smith *et al*. (7), and PC_A21 (9) from Smith *et al*. (7) The lung cancer data present in Recurrent Event is currently prepared in another manuscript.

### 1.4. Database

To make the scSVAS visualization more informative and user-friendly, we utilize several third-party databases and websites. Ensembl (10) database helps to locate the exact gene and transcript in CNV Heatmap, Clonal Lineage, Recurrent Event applications. UCSC genome browser (11) provides the cytobands, non-N region, and genome repeat annotation for visualization applications if applicable. In Cell Phylogeny, Clonal Lineage, and Recurrent Event, redirection to GeneCards (12) www.genecards.org is offered. Furthermore, MsigDB (v7.2) (13) database is adopted for gene set annotation in Clonal Lineage and Recurrent Event.

## Supplementary Note 2: Supplementary Manual

### 2.1. CNV View

Over the past two decades, CNV heatmap has often been adopted to visualize the CNV profiles of a batch of samples or single-cells in various sequencing protocols. e.g. bulk SNP array (1, 2), whole genome/exon sequencing (3), single-cell RNA sequencing (4–6). For single-cell DNA sequencing, CNV heatmap aids the landscape view of single-cell copy numbers in several literature pieces (7–11). In addition, Smith *et. al*. developed a visualization tool CellScape (packed in E-scape) for single-cell CNV heatmap (12).

Nevertheless, with the stride of 10x genomics high-throughput sequencing, the scale of sequenced cells escalates exponentially, aka, thousands of cells at a time (13–15). Efficient visualization of the heatmap with a large (e.g. 1k × 5k) size is critical for scientific interpretation. Plotting using R, Python packages, or existing heatmap visualization tools like CellScape are incredibly time-consuming and memory-intensive when it reaches thousands of cells and thousands of genomic regions.

It is essential to reduce the size of the heatmap while retaining the heterogeneity among single-cells. The 10x CNV official visualization tool Loupe (13) solves this issue by building a single-cell dendrogram in advance, splitting single-cells into less than 100 subgroups by cutting the dendrogram, and collapsing single-cells inside the cluster into one row in the heatmap. Cluster zoom-in/out operation is achieved by clicking the node in the dendrogram.

In the scSVAS platform, cooperating the scSVAS pipeline, we build a web interface CNV View for interactive and real-time visualization of CNV landscape of scDNA-Seq data with a zoomable dendrogram. Compared with Loupe, CNV View also visualizes the aggregate subgroup CNV heatmap and stairstep, which is commonly adopted in reputable publications (9, 10). CNV View enables users to create the landscape of single-cell CNV profiles in two straightforward steps as:

1. Open https://sc.deepomics.org/demo-project/analyses/view in Google browser, and upload cnv profile file *_cnv.csv, predefined meta file *_meta.csv.
2. *Optional* With cnv profile file *_cnv.csv, predefined meta file *_meta.csv as inputs, users can run scSVAS to get the cut dendrogram of single-cells, and upload it to compress the heatmap plot.

Users can get a single-cell CNV landscape view of scDNA-Seq data, as shown in Supplementary Fig. S1. With single-cells as rows and genomic regions as columns, the cell CNV heatmap exhibits the copy number of a specific single-cell across the entire genome. The cell meta-heat map will be displayed on the left if users provide single-cell meta information. Besides, the aggregate subgroup CNV heat map and stairstep will also be listed in the bottom layers. The user can select the subgroups to be aggregated in the Editor General Settings. If users offer the cut dendrogram file, a zoomable cut dendrogram will be displayed on the left. If the mouse hovers over cell CNV/meta heatmap, cut dendrogram, and stairstep, an interactive tooltip carrying its vital information will appear.

#### 2.1.1. Interactions

- Download An SVG file will be generated when you click the Download button. We offer two themes, dark and light. To switch to the light theme, please click the Light Theme button.
- Tooltips and Highlights When your cursor hovers over an Oviz component on the visualization panel, essential information about the component will show up in the tooltip, and related components will be highlighted. There are several major types of Oviz components in the CNV View application, and their tooltipping and highlighting interactions are:
  – Unit Component on the Cell CNV Heatmap The tooltip will display the genome position and copy number of a unit. The name of the corresponding leaf node in the cut dendrogram will also be shown. Furthermore, the genome position, the leaf node, and the range of leaf nodes will be highlighted (Supplementary Fig. S1A).
  – Cut Dendrogram Node The tooltip will display the name of the current node, the number of cells in it, the parent node of it, and the distance between it and the root node. Further, the subtree and the covered cell range of the current node will be highlighted (Supplementary Fig. S1B).
  – Cut Dendrogram Branch The tooltip will display the names of the associated parent and child nodes and their branch distance. The branch, the parent node, and the child node will be highlighted (Supplementary Fig. S1C).
  – Unit Component on the Cell Meta Heatmap The tooltip will display the cell ID and meta-label of a unit. (Supplementary Fig. S1D).
  – Unit Component of the Aggregate Subgroup CNV Heatmap The tooltip will display the genome position, the copy number, and the subgroup name. (Supplementary Fig. S1E).
  – Component on the Aggregate Subgroup CNV Stairstep The tooltip will display the genome position and the average copy number of cells for all subgroups(Supplementary Fig. S1F).
- Dendrogram Zooming When users click a node in the cut dendrogram, the selected node will be regarded as the temporary tree root, and a new sub-cut dendrogram will be rendered. The cell CNV heatmap and meta panel will also be updated to fit the current cell range. When you click the Back to Root button, the whole CNV view will return to its initial status. You may also utilize the left arrow and right arrow buttons to un-do and re-do zooming operations (Supplementary Fig. S1A-B).

#### 2.1.2. Editor Functionality

The editor offers various options to fine-tune the visualization. Users can adjust the editor width and font size in Editor Settings.

- Demo File Sets Users can select a demo file set for an instant preview.
- Files
  – Upload Users can upload and manage the input files. Note that the repeated file name will be warned and given a random postfix.
  – Choose Users can choose files uploaded previously.
  – File sets Users can save multiple files together as a file set, then decide to display one file set previously stored.
- General Settings
  – Autoload heatmap Checkbox for users to decide whether the cell CNV heatmap will automatically be loaded.
  – NA events (separated by comma) User can define the NA events in CNV csv file, the default, N/A,NA means empty space, string N/A, and string NA will be considered NA events by file parser.
  – Reorder cells by Users can reorder cells in the cell CNV/meta heatmap by meta labels in ascending or descending order. The default order is the cell ordered in the uploaded CNV csv file. Please note this functionality is effective only when on cut dendrogram JSON file supplied.
  – Aggregate subgroup Users can select categorical meta-labels to display in the aggregate subgroup CNV heatmap and stairstep. The average copy number value in the subgroup will be displayed.
- Select a categorical meta label Users can choose which categorical meta-label to show in the heatmap. Note that this functionality is effective only when on cut dendrogram JSON files supplied.
- Layout Settings
  – Basic
    * Figure margin - left Users can adjust the left margin of the figure.
    * Figure margin - top Users can adjust the top margin of the figure.
    * Genome zoom slider - height Users can adjust the height of the genome zoom slider.
    * Margin between CNV heatmap and genome zoom slider. Users can adjust the margin between CNV heatmap and genome zoom slider.
    * Margin between cut dendrogram and meta heatmap. Users can adjust the margin between the cut dendrogram and meta-heatmap.
  – CNV Heatmap
    * Unit height of CNV heatmap, unit width of CNV heatmap (integer recommended) Users can adjust the unit height and unit width of CNV heatmap. Unit refers to the smallest rendered SVG object in the heatmap. Please note that the heatmap unit height and unit width are recommended to set as an integer. The floating-point will make the heatmap transparent owing to subpixel rendering.
    * Chromosomes - height Users can adjust the height of chromosomes.
    * Show the vertical line between chromosomes User can decide whether to show vertical line between chromosomes.
    * Desired width of CNV heatmap. Users can adjust the width of CNV heatmap, default is 1000.
    * Left highlight - width User can adjust the width of left highlight.
    * Top highlight - height User can adjust the height of right highlight.
  – Meta Heatmap
    * Unit width of meta heatmap User can adjust the unit width of meta-heatmap.
    * Meta heatmap legend - width User can adjust the width of meta-heatmap legend.
    * Margin between different meta-heatmap legends User can adjust the margin between different meta-heatmap legends.
  – Cut Dendrogram
    * Dendrogram - width User can adjust the width of cut dendrogram.
- Color palettes Users can customize color palettes for available categorical meta labels and continuous meta labels.

### 2.2. CNV Heatmap

The web interface CNV Heatmap is an extended version of CNV View by supporting extra functionalities, including genome region zooming, local region search, and local region annotation.

CNV Heatmap enables users to create the cnv heatmap of single-cell CNV profiles in following straightforward steps as:

1. Open https://sc.deepomics.org/demo-project/analyses/heatmap in Google browser, and upload cnv profile file *_cnv.csv, predefined meta file *_meta.csv.
2. *Optional* With cnv profile file *_cnv.csv, predefined meta file *_meta.csv as inputs, users can run scSVAS to get the cut dendrogram of single-cells, and upload it to compress the heatmap plot.
3. *Optional* Users can also upload a customized bed file to check the CNV heatmap in local region.

Users can get a single-cell CNV landscape view of scDNA-Seq data, as shown in Supplementary Fig. S2A. With single-cells as rows and genomic regions as columns, the cell CNV heatmap exhibits the copy number of a specific single-cell across the entire genome. The cell meta-heat map will be displayed on the left if users provide single-cell meta information. Besides, there is a genome zoom slider located at the bottom of CNV heatmap. Users can drag the slider to zoom in and out on the genome region. Users can also upload a bed region file or search for a local genome region in Editor General Settings. If the local genome region is less than 5M, an annotation layer including Repeat track and Gene track will be displayed (Supplementary Fig. S2B). If the mouse hovers over the cell CNV/meta heatmap, repeats, and genes, an interactive tooltip carried its vital information will appear.

#### 2.2.1. Interactions

- Download An SVG file will be generated when you click the Download button. We offer two themes, dark and light. To switch to the light theme, please click the Light Theme button.
- Tooltips and Highlights When your cursor hovers over an Oviz component on the visualization panel, essential information about the component will show up in the tooltip, and related components will be highlighted. There are several major Oviz components in the CNV Heatmap application, and their tooltipping and highlighting interactions are:
  – Cell CNV Heatmap, Cell Meta Heatmap, Cut Dendrogram Node, Cut Dendrogram Branch Tooltipping and highlighting interactions are the same with CNV View visualization (Supplementary Fig. S2A, Supplementary Fig. S1A-D).
  – Gene Track Tooltip will display the transcript name and gene body interval for the covered gene. Tooltip will display exon number and exon intervals for covered gene exon. (Supplementary Fig. S2B).
  – Repeat Track Tooltip will display the names of repeat class, repeat element, repeat family, genome position, and strand information of a selected repeat (Supplementary Fig. S2B).
- Zoom Slider Button Users can click this button with active genome zoom slider mode.
- Bed Region Button Users can click this button to active bed region mode.
- Search Bed Button Users can click this button to active search bed mode.
- Genome Zoom Slider Users can adjust and drag the slider along the genome region to zoom in and out (Supplementary Fig. S2A).
- Local Region Zoom Slider Users can adjust and drag the slider along the local region to zoom in and out (Supplementary Fig. S2B).
- Pages Users can click Prev or Next to switch uploaded local bed regions.
- Dendrogram Zooming Dendrogram zooming is the same as CNV View visualization.

#### 2.2.2. Editor Functionalities

The editor offers various options to fine-tune the visualization. Users can adjust the editor width and font size in Editor Settings.

- Demo File Sets, Files Demo file sets and files Functionalities are the same with CNV Heatmap visualization.
- General Settings
- Auto load heatmap, NA events, Reorder cells by Auto load heatmap, NA events, Reorder cells by functionality are the same with CNV View visualization.
- Mode Users can switch from Zoom Slider, Bed Region, or Search Bed modes.
- Select categorical meta label, Layout Settings Select categorical meta-label, Layout Settings are the same with CNV View visualization.
- Search (Available in Bed Region and Search Bed mode)
  – Search local genome region Users can enter a local genome region to display, e.g. chr17:5,565,097-9,590,863. This functionality is only effective when the search bed mode is activated.
  – Bed region Users can select the local genome region in uploaded bed file to display. This functionality is only effective when bed region mode is activated.
- Genome Browser (Available in Bed Region and Search Bed mode)
  – Annotation Panel
    * Display strand Users can choose to display positive, negative, or both strands on the gene track.
    * Show repeat track Users can choose to display the repeat track or not.
    * Merge gene tracks Users can choose to merge gene track to get condensed displaying.
    * Display gene name Users can choose to display the gene name or transcript name in the gene track.
    * Display all genes Users can choose to display all genes on the gene track.
  – Genes This section lists all genes in the current local region, users can decide to display which transcript of one gene.
- Color palettes Users can customize color palettes for available categorical meta labels and continuous meta labels.

### 2.3. Cell Phylogeny

In scSVAS platform, we build a readily available web interface Cell Phylogeny for interactive and real-time visualization of scDNA-Seq data, focusing on the zoomable dendrogram.

Cell Phylogeny enables users to create the cell phylogeny plot in one straightforward step as:

1. With cnv profile file *_cnv.csv, predefined meta file *_meta.csv and targeted gene list as inputs, run scSVAS to get the single-cells cut dendrogram *_cut.json, the subclone and embedding results *_meta_scsvas.csv, and targeted gene cnv profiles *_gene_cnv.csv.
2. Open https://sc.deepomics.org/demo-project/analyses/cell_phylogeny in Google browser, and upload files *_cut.json, optional files *_meta_scsvas.csv and *_gene_cnv.csv.

Then, you may get a zoomable cut dendrogram of single-cells, as illustrated in Supplementary Fig. S3. Users can switch the tree between topdown and circular modes (Supplementary Fig. S3A-B). If the optional file *_meta.csv and *_gene_cnv.csv are uploaded, the cell meta and CNV heatmap (on gene or local bins) will be shown. Users can decide to display or hide these meta labels in the Editor-Select categorical meta label. If the mouse hovers over a dendrogram or heatmap, an interactive tooltip carried its vital information will appear.

#### 2.3.1. Interactions

- Download An SVG file will be generated when you click the Download button. We offer two themes, dark and light. To switch to the light theme, please click the Light Theme button.
- Tooltips and Highlights When your cursor hovers over an Oviz component on the visualization panel, essential information about the component will show up in the tooltip, and related components will be highlighted. There are several major types of Oviz components in the CNV View application, and their tooltipping and highlighting interactions are:
  – Cut Dendrogram Node, Cut Dendrogram Branch, Dendrogram Zooming, Meta CNV Heatmap Tooltipping and highlighting interactions are the same with CNV View visualization (Supplementary Fig. S1A-D).
  – Cell CNV Heatmap The tooltip will display the column name of a unit (such as gene or bin region) and its corresponding leaf node in the cut dendrogram. Furthermore, the column name, the leaf node, and the leaf node’s range will be highlighted (Supplementary Fig. S3A-B).
- TopDown <=> Circular Users can click this button to switch the cut dendrogram between TopDown and Circular modes (Supplementary Fig. S3A-B).

#### 2.3.2. Editor Functionalities

The editor offers various options to fine-tune the visualization. Users can adjust the editor width and font size in Editor Settings.

- Demo File Sets, Files Demo file sets and files Functionalities are the same with CNV View visualization.
- General Settings
  – Type of cut dendrogram Users can choose the type of cut dendrogram between topdown and circular.
  – NA events (seperated by comma,) User can define the NA events in CNV csv file, the default, N/A,NA means empty space, string N/A, and string NA will be considered NA events by file parser.
- Topdown Layout Settings Topdown Layout Settings are the same with Layout Settings in CNV Cell Phylogeny visualization.
- Circular Layout Settings
  – Basic
    * Figure margin - left Users can adjust the left margin of the figure.
    * Figure margin - top Users can adjust the top margin of the figure.
    * Margin between circular cut dendrogram and meta heatmap Users can adjust the margin between circular cut dendrogam and meta-heatmap.
  – Circular Layout
    * Start angle of circular cut dendrogam Users can adjust the start angle of circular cut dendrogram.
    * End angle of circular cut dendrogam Users can adjust the end angle of circular cut dendrogram.
    * Inner radius of circular cut dendrogram Users can adjust the inner radius of circular cut dendrogram.
    * Unit width of CNV heatmap (integer recommended) Users can adjust the unit width of CNV and meta heatmap. Unit refers to the smallest rendered SVGobject in heatmap. Please note that the heatmap unit width are recommend to set as integer, floating point will make heatmap transparent owing to subpixel rendering.
- Color Palettes Users can customize color palettes for available categorical meta labels and continuous meta labels.

### 2.4. Ploidy Stairstep

As previously mentioned, for single-cell DNA cancer data, the ploidy distribution can intuitively show tumor heterogeneity. The ploidy line plot along the chromosomes can also visually show the heterogeneity between tumor subclones by combining genomic coordinates. By collapsing the single-cells in the same tumor subclones into one observation, we can infer each subclone’s pseudo-bulk ploidy. Since cancer CNV ploidy line alters along chromosomes, we call it the stairstep plot.

In scSVAS platform, we develop a readily available web interface Ploidy Stairstep for interactive and real-time visualization of ploidy stairstep plot for scDNA-Seq data.

Ploidy Stairstep enables users to create the ploidy stairstep plot just in one simple step as:

1. Open https://sc.deepomics.org/demo-project/analyses/ploidy_stairstep in Google browser, and upload cnv profile file *_cnv.csv and predefined meta file *_meta.csv.
2. *Optional* With cnv profile file *_cnv.csv and predefined meta file *_meta.csv as inputs, users may also run scSVAS to get the subclone cluster result in *_meta_scsvas.csv first.

Then, you may get a matrix of the ploidy stairstep plot of scDNA-Seq data, as illustrated in Supplementary Fig. S4. The columns will list all categorical meta labels in the uploaded file *_meta_scsvas.csv by default. The first row exhibits all subgroups’ ploidy stairstep for specific categorical meta labels in an aggregate form. The second line displays the ploidy stairstep of pseudo-bulk profiles. Then, the following rows will list the ploidy stairstep of all available subsets individually. Users can decide to display or hide these meta labels and subsets in the Editor-Select categorical meta label. If the mouse hovers over the stairstep plot, an interactive tooltip carried its vital information will appear.

#### 2.4.1. Interactions

- Download An SVG file will be generated when you click the Download button. We offer two themes, dark and light. To switch to the light theme, please click the Light Theme button.
- Tooltips and Highlights When your cursor hovers over an Oviz component on the visualization panel, essential information about the component will show up in the tooltip, and related components will be highlighted. There are two major types of Oviz components in the CNV Ploidy Stairstep application and their tooltipping and highlighting interactions are:
  – Stairstep plot The tooltip will display the genome position and the average copy number (Supplementary Fig. S4).
  – Aggregate subgroup distribution plot The tooltip will display the genome position and the average copy number for each subgroup respectively (Supplementary Fig. S4).

#### 2.4.2. Editor Functionalities

The editor offers various options to fine-tune the visualization. Users can adjust the editor width and font size in Editor Settings.

- Demo File Sets, Files Demo file sets and files Functionalities are the same with CNV View visualization.
- General Settings
  – Maximum CN value Users can adjust the maximum CN value of y-axis.
  – Stairstep plot height Users can adjust the height of each stairstep plot.
  – Stairstep plot width Users can adjust the width of each stairstep plot.
  – Stairstep plot line width Users can adjust the line width of each stairstep plot.
- Select categorical meta label Users can choose which categorical meta labels to display.
- Color Palettes Users can customize color palettes for available categorical labels.

### 2.5. Ploidy Distribution

Over the past decade, the single-cell DNA CNV calling algorithms usually model the relationship between sequenced read count and copy number employing Poisson distribution with mappability, GC content, scale factor as confounding factor (11, 13, 16, 17).

In cancer datasets, the copy number of different regions may encounter dramatic gain or loss (18–21). For single-cell DNA cancer data, ploidy distribution can intuitively display the tumor intra-heterogeneity.

To address this concern, in scSVAS platform, we develop a readily available web interface Ploidy Distribution for interactive and real-time visualization of ploidy density plots for scDNA-Seq data.

Ploidy Distribution enables users to create the ploidy density plot just in one simple step as:

1. Open https://sc.deepomics.org/demo-project/analyses/ploidy_distribution in Google browser, and upload cnv profile file *_cnv.csv and predefined meta file *_meta.csv.
2. *Optional* With cnv profile file *_cnv.csv and predefined meta file *_meta.csv as inputs, users may also run scSVAS to get the subclone cluster result in *_meta_scsvas.csv first.

Then, you may get a matrix of the ploidy density plot of scDNA-Seq data, as illustrated in Supplementary Fig. S5. The columns will list all categorical meta labels in the uploaded file *_meta_scsvas.csv by default. The first row exhibits all subgroups’ ploidy distribution for specific categorical meta labels in an aggregate form. The second line displays the ploidy distribution of total single-cells, just like the way for bulk sequencing. Then, the following rows will list the ploidy distribution of all available subsets individually. Users can decide to display or hide these meta labels and subsets in the Editor-Select categorical meta label. Users can choose to set the ploidy unit as cell mean ploidy or bin ploidy in Editor-Mode. If the mouse hovers over the density plot, an interactive tooltip carried its vital information will appear.

#### 2.5.1. Interactions

- Download An SVG file will be generated when you click the Download button. We offer two themes, dark and light. To switch to the light theme, please click the Light Theme button.
- Tooltips and Highlights When your cursor hovers over an Oviz component on the visualization panel, essential information about the component will show up in the tooltip, and related components will be highlighted. There are two major types of Oviz component in the CNV Ploidy Distribution application and their tooltipping and highlighting interactions are:
  – Distribution plot The tooltip will display the current ploidy value and the density of that ploidy. (Supplementary Fig. S5).
  – Aggregate subgroup distribution plot The tooltip will display the current ploidy value and the density of that ploidy for each subgroup. (Supplementary Fig. S5).

#### 2.5.2. Editor Functionalities

The editor offers various options to fine-tune the visualization. Users can adjust the editor width and font size in Editor Settings.

- Demo File Sets, Files Demo file sets and files Functionalities are the same with CNV View visualization.
- General Settings
  – Mode Users can select to display cell mean ploidy or bin ploidy mode.
  – Get log10-scale of count? Users can choose whether to apply log10-scale on density count.
  – Set total number of intervals (bins) Users can set the total number of bins. The bin is the block that you use to combine values before getting the frequency.
  – Only get the plot with 10+ data? Users can decide whether filter the subgroup plot with less than 10 data points.
  – Distribution plot height Users can adjust the height of each distribution plot.
  – Distribution plot width Users can adjust the width of each distribution plot.
  – Maximum value of ploidy Users can adjust the maximum value of ploidy in the x-axis.
- Select categorical meta label Users can choose which categorical meta labels to display.
- Color Palettes Users can customize color palettes for available categorical meta labels.

### 2.6. Embedding Map

High-dimensional data could be challenging to visualize. Reducing data into two dimensions is essential for representing the inherent structure of the data. Several supervised and unsupervised embedding methods have been proposed and widely applied to multiple disciplines in the past two decades. For example, linear dimension reduction tools like Principal Component Analysis (PCA), Independent Component Analysis (ICA) (22), Non-negative Matrix Factorization (NMF) (23) specify distinct rubrics to conduct linear projection of data. Furthermore, to tackle nonlinear data structure, t-distributed Stochastic Neighbor Embedding (t-SNE) (24), Uniform Manifold Approximation and Projection (UMAP) (25), and Potential of Heat-diffusion for Affinity-based Trajectory Embedding (PHATE) (26) are developed. We also build a Matrix Factorization-based Deep neural network (DeepMF) (27), which is compatible with both linear and non-linear embedding.

In scSVAS platform, we build a readily available web interface Embedding Map for interactive and real-time visualization of scDNA-Seq data.

With the advance of sing-cell DNA sequencing techniques, the CNV data of tens of thousands of cells could be profiled simultaneously. In terms of large-scale data, conventional 2D scatter plots may disguise essential information. Embedding Map defeats this overplotting obstacle utilizing hexagonal binning (28, 29), which also has benefits on time and memory complexity.

Embedding Map enables users to create the 2D embedding plot in two straightforward steps as:

1. With cnv profile file *_cnv.csv, predefined meta file *_meta.csv and targeted gene list as inputs, run scSVAS to get the 2D embedding (PCA, ICA, TSNE, UMAP, PHATE, and DeepMF) results *_meta_scsvas.csv and targeted gene cnv profiles *_gene_cnv.csv.
2. Open https://sc.deepomics.org/demo-project/analyses/embedding_map in Google browser, and upload files *_meta_scsvas.csv and *_gene_cnv.csv.

Then, you may get a matrix of 2D representations of scDNA-Seq data, as illustrated in Supplementary Fig. S6A. The column will list all embedding techniques available in the uploaded file *_meta_scsvas.csv by default. Users can decide to display or hide these embedding methods in the Editor-General Settings. The rows show different strategies to color the single-cell data point. If the hexagon mode is activated, the embedded cells colored with density will be displayed (Supplementary Fig. S6B). If the optional file *_gene_cnv.csv is uploaded, the embedded cells colored with gene CNV profiles will be shown. Users can specify the gene for coloring in the Editor-General Settings. Moreover, all categorical meta labels available in the uploaded file *_meta_scsvas.csv will be used as color schemes by default. Users can decide to display or hide these meta labels in the Editor-Select categorical meta label. If the mouse hovers over one scatter point or hexagon bin, an interactive tooltip carried its vital information will appear.

#### 2.6.1. Interactions

- Download An SVG file will be generated when you click the Download button. We offer two themes, dark and light. To switch to the light theme, please click the Light Theme button.
- Tooltips and Highlights When your cursor hovers over an Oviz component on the visualization panel, essential information about the component will show up in the tooltip, and related components will be highlighted. There are two major types of Oviz component in the CNV Embedding Map application and their tooltipping and highlighting interactions are:
  – Scatter point on the 2D-Embedding scatter plot The tooltip will display the x and y coordinates, the coloring value, and the cell ID. (Supplementary Fig. S6A).
  – Hexagon bins on the 2D-Embedding hexagon plot The tooltip will display the x and y coordinates, the coloring value, the number of cells in the bin, and the list of cell IDs (Supplementary Fig. S6B).

#### 2.6.2. Editor Functionalities

The editor offers various options to fine-tune the visualization. Users can adjust the editor width and font size in Editor Settings.

- Demo File Sets, Files Demo file sets and files Functionalities are the same with CNV View visualization.
- General Settings
  – Select embedding methods Users can select to display the available embedding methods.
  – Hexagon Mode Users can choose hexagon mode or scatter mode.
  – Width of hexagon bin Users can adjust the width of the hexagon bin.
  – Hexagon bin averaging scheme Users can choose mean or median as the hexagon bin averaging scheme.
  – Radius of scatter point Users can adjust the radius of the scatter point.
  – Search for gene Users can search or select a gene and color the embedding plot with its copy number.
  – Embedding plot height Users can adjust the height of each embedding plot.
  – Embedding plot width Users can adjust the width of each embedding plot.
- Select categorical meta label Users can choose which categorical meta label to use for coloring the embedded plot.
- Color Palettes Users can customize color palettes for Density and available categorical labels.

### 2.7. Time Lineage

Many studies have observed that intra-tumor heterogeneity (ITH) is one of the principal causes of cancer therapy-resistant, tumor recurrence, and deaths (30). Over the past decades, researchers are interested in studying the clonal dynamics from multiple timepoint. For example, the time lineage between subclones before and after therapeutic intervention (10, 31).

Herein, in scSVAS platform, we develop a readily available web interface Time Lineage for interactive and real-time visualization of clonal lineage over time for scDNA-Seq data.

Time Lineage enables users to create the time lineage visualization just in following steps:

1. Open https://sc.deepomics.org/demo-project/analyses/time_lineage in Google browser, and upload the customized clonal tree file clonal_edges.csv and clonal prevalence file clonal_prev.csv.

Time Lineage is composed of fishplot, lineage tree, and cellular ensemble (Supplementary Fig. S7A). Fishplot conceptually manifests the proportion of tumor subclones at different tumorigenesis stages along time. We use the bezier curve to fit the trend of subclones over time. Two distinct head shapes (bullet and onion) and three layouts (stack, space, and center) are offered (Supplementary Fig. S7B). The lineage tree exhibits the evolutionary relationship between tumor subclones. Users can choose different tree shapes from topdown normal, circular normal, and circular acute (Supplementary Fig. S7C). The cellular ensemble is an abstract aesthetic presenting the tumor’s cellular prevalence at a certain point in time (Supplementary Fig. S7D). If the mouse hovers over the subclone in fishplot, lineage tree, and cellular ensemble, an interactive tooltip carried its vital information will appear (Supplementary Fig. S7).

#### 2.7.1. Interactions

- Download An SVG file will be generated when you click the Download button. We offer two themes, dark and light. To switch to the light theme, please click the Light Theme button.
- Tooltips and Highlights When your cursor hovers over an Oviz component on the visualization panel, essential information about the component will show up in the tooltip, and related components will be highlighted. There are several major types of Oviz component in the CNV Time Lineage application and their tooltipping and highlighting interactions are:
  – Subclone in fishplot The tooltip will display the name of the subclone, the clone prevalence at each timepoint. The corresponding subclone in fishplot and lineage tree will be highlighted (Supplementary Fig. S7A-B).
  – Lineage tree branch The tooltip will display the parent node name, child node name, and branch’s distance. The corresponding branch and nodes will be highlighted in fishplot and cellular ensemble (Supplementary Fig. S7A).
  – Lineage tree node The tooltip will display the subclone node name, distance to the root node, clonal frequency, the number of cells in the subclone. The corresponding subclone in fishplot and cellular ensemble will be highlighted (Supplementary Fig. S7C).
  – Cellular ensemble The tooltip will display the name and prevalence of the clone. The corresponding clones will be highlighted in lineage tree and fishplot (Supplementary Fig. S7D).

#### 2.7.2. Editor Functionalities

The editor offers various options to fine-tune the visualization. Users can adjust the editor width and font size in Editor Settings.

- Demo File Sets, Files Demo file sets and files Functionalities are the same with CNV View visualization.
- General Settings
  – Fishplot
    * Show clone name Users can decide show clone name or not.
    * X-axis title of fishplot Users can change the title for x-axis, default is Time Point.
    * Y-axis title of fishplot Users can change the title for y-axis, default is Clonal Prevalence.
    * Vertical layout of subclones Users can select the vertical layout of subclones from stack, space, and center.
    * Head shape of fishplot Users can select the shape of clone head from bullet or onion.
    * Width of fishplot Users can set the width of fishplot.
    * Height of fishplot Users can set the height of fishplot.
  – Lineage Tree
    * Title of lineage tree Users can change the title for lineage tree. default is Clonal Lineage Tree.
    * Type of lineage tree Users can change the type of lineage tree.
  – Cellular Ensemble
    * Number of cells Users can adjust the number of cells in cellular ensemble presentation.
    * Radius of cellular aggregation Users can adjust the radius of cellular ensemble presentation.
- Color Palettes Users can customize color palettes for subclones.

### 2.8. Space Lineage

In scSVAS platform, we develop a readily available web interface Space Lineage for interactive and real-time visualization of clonal lineage across time for scDNA-Seq data.

Space Lineage enables users to create the clonal and spatial lineage visualization just in following steps:

1. Open https://sc.deepomics.org/demo-project/analyses/space_lineage in Google browser, and upload files subclone tree file clona_ledges.csv, clonal prevelance file clonal_prev.csv.
2. *Optional* With subclone tree file clonal_edges.csv and clonal prevelance file clonal_prev.csv as input, run scSVAS space.py to get the space tree results space_edges.csv, then upload it.
3. *Optional* Users can also upload customized lesion images.

Space Lineage displays the clonal dynamics across subclones and lesions (Supplementary Fig. S8A). We offer subclone and lesion lineage trees. Supplementary Fig. S8B-C provide a zoomable human anatomy atlas (license-free). Users can select different anatomy images from cartoon anatomy atlas (male, female, brain, intestines, kidney, liver, lung, pancreas, stomach, thymus, thyroid, urinary system, male reproductive system, and female reproductive system). Instead of the arduous lesion position precalculation required in E-Scape (12), we provide an image pointer for each lesion. Users can right-click to active it and drag the pointer to the exact lesion position (Supplementary Fig. S8D). If the mouse hovers over the branch and node in the lineage tree, an interactive tooltip carried its vital information will appear (Supplementary Fig. S8A).

#### 2.8.1. Interactions

- Download An SVG file will be generated when you click the Download button. We offer two themes, dark and light. To switch to the light theme, please click the Light Theme button.
- Tooltips and Highlights When your cursor hovers over an Oviz component on the visualization panel, essential information about the component will show up in the tooltip, and related components will be highlighted. There are several major types of Oviz component in the Space Lineage application and their tooltipping and highlighting interactions are:
  – Subclone lineage tree node The tooltip will display the name of subclone/lesion, the clone prevalence at each timepoint if available. The corresponding subclone/lesion in lineage tree, fishplot will be highlighted (Supplementary Fig. S8A).
  – Lesion lineage tree node The tooltip will display the lesion node name, distance to the root node, clonal frequency, the number of cells in the subclone. The corresponding lesion in other lineage trees will be highlighted. (Supplementary Fig. S8A).
  – Lineage tree branch The tooltip will display the parent node name, child node name, and branch’s distance. The corresponding branch and nodes will be highlighted in other lineage trees (Supplementary Fig. S8A).
- Movable anatomy circle We provide a movable image pointer. Users can right-click the image circle to active and deactivate it. When the image circle is activated, users can drag the circle to anywhere they want in the anatomy image (Supplementary Fig. S8C).
- Anatomy image pointer For each lesion, we provide an image pointer. Users can right-click the image pointer to active and deactivate it. When the image pointer is activated, users can drag the pointer to anywhere they want in the anatomy image (Supplementary Fig. S8D).

#### 2.8.2. Editor Functionalities

The editor offers various options to fine-tune the visualization. Users can adjust the editor width and font size in Editor Settings.

- Demo File Sets, Files Demo file sets and files Functionalities are the same with CNV View visualization.
- General Settings
  – Lineage Tree
    * Show subclone name Users can decide show clone name or not.
    * Type of lineage tree Users can change the type of lineage tree.
    * Show low prevalence clone Users can decide whether to show low prevalence clone or not.
    * Low prevalence threshold Users can change the value of low prevalence threshold, default it 0.01.
  – Anatomy type Users can select different anatomy images from cartoon anatomy atlas (Supplementary Fig. S8B).
  – Radius of anatomy image Users can adjust the radius of anatomy image (Supplementary Fig. S8C).

### 2.9. Space Prevalence

In scSVAS platform, we develop a readily available web interface Space Prevalence for interactive and real-time visualization of clonal dynamics across multiple samples.

Space Prevalence enables users to create the space prevalence visualization just in following steps:

1. With subclone tree file clonal_edges.csv and clonal prevalence file clonal_prev.csv as input, run scSVAS space.py to get the space tree results space_edges.csv.
2. Open https://sc.deepomics.org/demo-project/analyses/space_prevalence in Google browser, and upload the customized clonal tree file clonal_edges.csv, clonal prevalence file clonal_prev.csv and space tree results space_edges.csv.

Space Prevalence displays the clonal tree, lesion tree, and space prevalence across subclones and lesions. If the mouse hovers over the node in the lineage tree and Sankey diagram, an interactive tooltip carried its vital information will appear (Supplementary Fig. S8).

#### 2.9.1. Interactions

- Download An SVG file will be generated when you click the Download button. We offer two themes, dark and light. To switch to the light theme, please click the Light Theme button.
- Tooltips and Highlights When your cursor hovers over an Oviz component on the visualization panel, essential information about the component will show up in the tooltip, and related components will be highlighted. There are several major types of Oviz component in the Space Prevalence application and their tooltipping and highlighting interactions are:
  – Subclone lineage tree node The tooltip will display the name of the subclone/lesion, the clone prevalence at each timepoint if available. The corresponding subclone/lesion in lineage tree, fishplot, and Sankey plot will be highlighted (Supplementary Fig. S9).
  – Lesion lineage tree node The tooltip will display the lesion node name, distance to the root node, clonal frequency, the number of cells in the subclone. The corresponding lesion in the Sankey plot will be highlighted (Supplementary Fig. S9).
  – Lineage tree branch The tooltip will display the parent node name, child node name, and branch’s distance. The corresponding branch and nodes will be highlighted in the Sankey plot and clonal prevalence matrix (Supplementary Fig. S9).
  – Sankey plot node The tooltip will display the subclone/lesion node name. The corresponding subclone/lesion in lineage tree will be highlighted (Supplementary Fig. S9).
  – Sankey plot link The tooltip will display the subclone node name, lesion node name, and the clonal prevalence across them. The corresponding branch and nodes will be highlighted in the lineage tree and clonal prevalence matrix (Supplementary Fig. S9).
  – Clonal prevalence matrix tile The tooltip will display the subclone node name, lesion node name, and the clonal prevalence across them. The corresponding branch and nodes will be highlighted in the lineage tree and Sankey plot (Supplementary Fig. S9).

#### 2.9.2. Editor Functionalities

The editor offers various options to fine-tune the visualization. Users can adjust the editor width and font size in Editor Settings.

- Demo File Sets, Files Demo file sets and files Functionalities are the same with CNV View visualization.
- General Settings
  – Lineage Tree
    * Show subclone name Users can decide show clone name or not.
    * Type of lineage tree Users can change the type of lineage tree.
    * Show low prevelance clone Users can decide whether to show low prevalence clone or not.
    * Low prevalence threshold Users can change the value of low prevalence threshold, default it 0.01.

### 2.10. Clonal Lineage

Many studies have observed that intra-tumor heterogeneity (ITH) is one of the principal causes of cancer therapy-resistant, tumor recurrence, and deaths (30). An accurate understanding of the subclone structure and evolutionary history benefits precise treatments for individual patients (32). Over the past decades tools utilize SNV (33), CNV information (34), or combine these two phenotype markers (35) to infer the phylogeny tree. There are tools customized for different sequencing protocols, including multi-region (36), single-cell (37–39).

Unlike the traditional phylogenetic trees as visualized in Cell Phylogeny, we focus on the clonal lineage tree, which more accurately reflects the process of tumor evolution. In a clonal lineage tree, ancestors and offspring tumor cells/subpopulations can coexist simultaneously; therefore, the internal node can be the single-cell/subpopulation we observed. The tumor accumulates mutations over evolution time, and child tumor cell/subpopulation carries parental and newly-acquired aberrations. The tree linkage between parent and child node is more about asymmetric subset connections than symmetric distance.

There are several tools to visualize the clonal lineage tree with subclones as tree nodes. For example, fishplot presents clonal dynamics over time (40–42); sphere of cells present clonal subpopulations of a sample (12, 42), and annotated node-based (12, 42) and branch-based trees present clonal relationships and seeding patterns between samples (42). Nevertheless, there is no good tool to display the acquired mutations over time.

To address this concern, in scSVAS platform, we develop a readily available web interface Clonal Lineage for interactive and real-time visualization of clonal lineage and associated CNV across time for scDNA-Seq data.

Clonal Lineage enables users to create the clonal lineage visualization just in following steps:

1. With cnv profile file *_cnv.csv, predefined meta file *_meta.csv and targeted gene list as inputs, run scSVAS to get the build clonal lineage results *_evo.json.
2. Open https://sc.deepomics.org/demo-project/analyses/clonal_lineage in Google browser, and upload the customized clonal tree file *_evo.json.
3. *Optional* Users can also upload the predefined gene list target_anno.tsv to only display the CNV shift of targeted gene.

Clonal Lineage comprises lineage tree, group CNV heatmap, cellular ensemble, lineage tree branch, stairstep, and gene box. The lineage tree exhibits the evolutionary relationship between tumor subclones (Supplementary Fig. S10A). Users can choose different tree shapes from topdown normal, circular normal, circular acute, and fishplot (Supplementary Fig. S10B). Fishplot conceptually manifests the proportion of tumor subclones at different tumorigenesis stages over time. We use the bezier curve to fit the trend of subclones over time. Two distinct head shapes (bullet and onion) are offered (Supplementary Fig. S7B). The cellular ensemble is an abstract aesthetic presenting the tumor’s cellular prevalence at a certain time point (Supplementary Fig. S10A). The group CNV heatmap displays the averaged copy number profiles of subclones (Supplementary Fig. S10A). The lineage tree branch displays the number of gain and loss regions for each tree branch (Supplementary Fig. S10A). The stairstep and gene box depict the detailed CNV shift from the parent node to the child node (Supplementary Fig. S10A). Users can click the lineage tree branch to check the different CNV shifts. If the mouse hovers over lineage tree, group CNV heatmap, cellular ensemble, lineage tree branch, stairstep, and gene box, an interactive tooltip carried its vital information will appear (Supplementary Fig. S10A).

#### 2.10.1. Interactions

- Download An SVG file will be generated when you click the Download button. We offer two themes, dark and light. To switch to the light theme, please click the Light Theme button.
- Tooltips and Highlights When your cursor hovers over an Oviz component on the visualization panel, essential information about the component will show up in the tooltip, and related components will be highlighted. There are several major types of Oviz component in the Clonal Lineage application and their tooltipping and highlighting interactions are:
  – Lineage tree node The tooltip will display the subclone node name, distance to the root node, clonal frequency, the number of cells in the subclone. The corresponding node in aggregate subclone CNV heatmap, lineage tree branch will be highlighted (Supplementary Fig. S10A).
  – Lineage tree branch The tooltip will display the parent node name, child node name, and branch’s distance. The corresponding branch will be highlighted (Supplementary Fig. S10A).
  – Subclone in fishplot The tooltip will display the name of subclone, the clone prevalence at each timepoint. The corresponding subclone in fishplot, lineage tree, and aggregate subclone CNV heatmap will be highlighted. (Supplementary Fig. S10A).
  – Aggregate subclone CNV heatmap The tooltipping and highlighting interactions are the same with aggregate subgroup CNV heatmap in CNV View application (Supplementary Fig. S1E).
  – Cellular ensemble The tooltip will display the name and prevalence of the clone. The corresponding clones will be highlighted in lineage tree or fishplot (Supplementary Fig. S10A).
  – Subclone branch The corresponding branch will be highlighted in lineage tree or fishplot (Supplementary Fig. S10A).
  – CNV shift Stairstep The tooltipping and highlighting interactions are the same with stairstep in Ploidy Stairstep application (Supplementary Fig. S4).
  – Gene box The corresponding gene box, cytoband, genomic position on stairstep will be highlighted (Supplementary Fig. S10A).
  – MsigDB Pathway The tooltip will display the name of the selected MsigDB pathway. The corresponding gene box, cytoband, genomic position on stairstep will be highlighted (Supplementary Fig. S10A).
  – Self-defined gene set. The tooltip will display the name of the self-defined gene set. The corresponding gene box, cytoband, genomic position on stairstep will be highlighted (Supplementary Fig. S10A).
- Subclone branch Users can display the CNV shifts of a particular subclone branch by clicking it.
- Packed gene box Users can click the packed gene box to look at the whole list of genes.
- External link on gene Users can jump to the GeneCards webpage by clicking on the gene listed in the gene box.
- External link on MsigDB pathway Users can jump to the MsigDB pathway webpage by clicking on the gene set icon.

#### 2.10.2. Editor Functionalities

The editor offers various options to fine-tune the visualization. Users can adjust the editor width and font size in Editor Settings.

- Demo File Sets, Files Demo file sets and files Functionalities are the same with CNV View visualization.
- General Settings
  – Basic
    * Select subclone label Users can select which subclone label to display.
  – Lineage Tree/Fishplot
    * Type of lineage tree Users can select the type of lineage tree from topdown normal, circular normal, circular acute, or fishplot (Supplementary Fig. S10B).
    * Vertical layout of subclones Users can select the vertical layout of subclones from stack, space, and center.
    * Head shape of fishplot Users can select the shape of clone head from bullet or onion.
  – Gene Box
    * CNV shift shreshold Users can adjust the CNV shift threshold. Only genes surpass this threshold will be displayed on gene box.
    * Maximum genes to display on gene box Users can adjust the maximum genes to display on gene box.
    * Aggregate gene sets Users can choose to aggregate the gene sets icon or not (Supplementary Fig. S10C).
- Layout Settings
  – Basic
    * Height of lineage tree/fishplot/aggregate subclone CNV heatmap Users can set the height of lineage tree/fishplot/aggregate subclone CNV heatmap.
    * Margin between aggregate subclone CNV heatmap and subclone branch Users can adjust the margin between aggregate subclone CNV heatmap and subclone branch.
    * Margin between subclone branch and CNV shift stairstep Users can adjust the margin between subclone branch and CNV shift stairstep.
    * Margin between CNV shift stairstep and legend Users can adjust the margin between CNV shift stairstep and legend.
  – Aggregate subclone CNV heatmap Aggregate subclone CNV heatmap functionalities are the same with CNV heatmap in CNV View visualization.
  – Subclone branch
    * Width of subclone branch Users can adjust the width of subclone branch.
    * Height of subclone branch Users can adust the height of subclone branch.
  – CNV shift stairstep
    * Maximum CN value Users can adjust the maximum CN value in y-axis.
    * Width of stairstep Users can adjust the width of the stairstep.
    * Height of stairstep Users can adjust the height of the stairstep.
    * Width of stairstep line Users can adjust the width of line in stairstep plot.
    * Margin between parent stairstep and child stairstep Users can adjust the margin between parent stairstep and child stairstep.
  – Gene Box
    * Width of gene box Users can adjust the width of gene box.
    * Column margin between gene boxes Users can adjust the margin between column margin between gene boxes.
- Gene sets selection Users can select the gene set to display by clicking on the checkbox.
- Color Palettes Users can customize color palettes for subclones, amp/loss, and gene sets.

### 2.11. Recurrent Event

In scSVAS platform, we also develop a readily available web interface Recurrent Event for interactive and real-time visualization of CNV profiles across multiple samples for scDNA-Seq data.

Recurrent Event enables users to create the recurrent event visualization just in following steps:

1. With group cnv profile file of multiple samples *_hcluster_cnv.csv as input, run scSVAS recurrent.py to get the recurrent event results *_recurrent.json.
2. Open https://sc.deepomics.org/demo-project/analyses/recurrent_event in Google browser, and upload the recurrent event results *_recurrent.json.
3. *Optional* Users can also upload the predefined gene list target_anno.tsv to only display the CNV shift of targeted gene.

Recurrent Event displays the CNV stairsteps of all samples (Supplementary Fig. **??**). All focal gains and losses are colored in red and blue by default. The gene box shows the recurrent genes and gene set annotation. If the mouse hovers over the subclone in stairstep and gene box, an interactive tooltip carried its vital information will appear.

#### 2.11.1. Interactions

- Download An SVG file will be generated when you click the Download button. We offer two themes, dark and light. To switch to the light theme, please click the Light Theme button.
- Tooltips and Highlights When your cursor hovers over an Oviz component on the visualization panel, essential information about the component will show up in the tooltip, and related components will be highlighted. There are several major types of Oviz component in the CNV Clonal Lineage application and their tooltipping and highlighting interactions are:
  – Stairstep The tooltipping and highlighting interactions are the same as stairstep in Ploidy Stairstep application.
  – Gene box The corresponding gene box, cytoband, genomic position on stairstep will be highlighted.
  – MsigDB gene sets The tooltip will display the name of selected MsigDB gene sets. The corresponding gene box, cytoband, genomic position on stairstep will be highlighted.
  – Self-defined gene set. The tooltip will display the name of the self-defined gene set. The corresponding gene box, cytoband, genomic position on stairstep will be highlighted.
- Packed gene box, External link on the gene, External link on MsigDB pathway The interaction is described in Clonal Lineage application.

#### 2.11.2. Editor Functionalities

The editor offers various options to fine-tune the visualization. Users can adjust the editor width and font size in Editor Settings.

- Demo File Sets, Files Demo file sets and files Functionalities are the same with CNV View visualization.
- General Settings
  – Maximum genes to display on gene box Users can adjust the maximum number of genes to display on the gene box, if the number of genes in one box surpasses the threshold, genes will be hidden.
  – Gap between samples Users can adjust the gap between samples.
  – Width of stairstep Users can adjust the width of the stairstep.
  – Height of stairstep Users can adjust the height of the stairstep.
- Gene sets selection Users can select which gene set to display.
- Sample and subclone selection Users can select which sample and subclone to display.

---

**Table S1.**
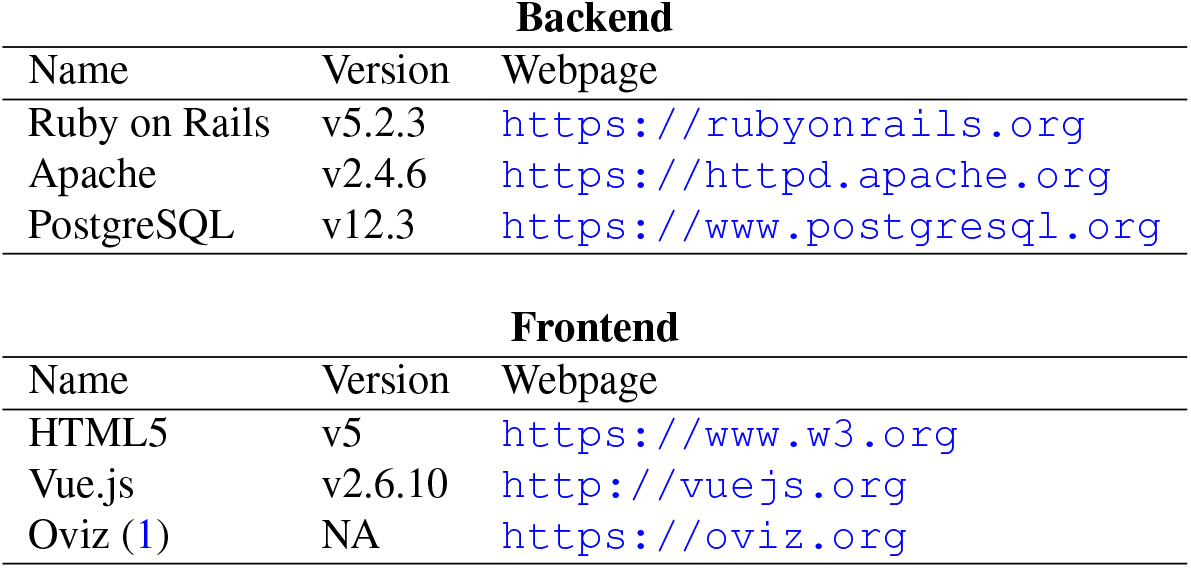
The technology stack of scSVAS. NA: Not Applicable.

| Backend |  |  |
| --- | --- | --- |
| Name | Version | Webpage |
| Ruby on Rails | v5.2.3 | https://rubyonrails.org |
| Apache | v2.4.6 | https://httpd.apache.org |
| PostgreSQL | v12.3 | https://www.postgresql.org |

**Table S1.** The technology stack of scSVAS. NA: Not Applicable.
| Frontend |  |  |
| --- | --- | --- |
| Name | Version | Webpage |
| HTML5 | v5 | https://www.w3.org |
| Vue.js | v2.6.10 | http://vuejs.org |
| Oviz (1) | NA | https://oviz.org |

## Supplementary Note 3: Supplementary Tables

**Table S2.** List of scSVAS applications’ demo datasets. Aberrations: Triple-negative breast cancer (TNBC); Acute myeloid leukaemia (AML); High grade serous ovarian cancer (HGSOC); Prostate cancer (PC); Lung cancer (LC).

| scSVAS Apps | Demo datasets | Description |
| --- | --- | --- |
| CNV View | TNBC_T10,T16 (2, 3), 10x_COLO829 (4) | TNBC_T10,T16: raw data obtained from Navin *et al.* (2), processed by SCYN (3) and scSVAS offline pipeline. 10x_COLO829: raw data obtained from Velazquez *et al.* (4) and processed by scSVAS offline pipeline. |
| CNV Heatmap | TNBC_T10,T16 (2, 3), 10x_COLO829 (4) | TNBC_T10,T16: raw data obtained from Navin *et al.* (2), processed by SCYN (3) and scSVAS offline pipeline. 10x_COLO829: raw data obtained from Velazquez *et al.* (4) and processed by scSVAS offline pipeline. |
| Cell Phylogeny | TNBC_T10,T16 (2, 3), 10x_COLO829 (4) | TNBC_T10,T16: raw data obtained from Navin *et al.* (2), processed by SCYN (3) and scSVAS offline pipeline. 10x_COLO829: raw data obtained from Velazquez *et al.* (4) and processed by scSVAS offline pipeline. |
| Ploidy Stairstep | TNBC_T10,T16 (2, 3), 10x_COLO829 (4) | TNBC_T10,T16: raw data obtained from Navin *et al.* (2), processed by SCYN (3) and scSVAS offline pipeline. 10x_COLO829: raw data obtained from Velazquez *et al.* (4) and processed by scSVAS offline pipeline. |
| Ploidy Distribution | TNBC_T10,T16 (2, 3), 10x_COLO829 (4) | TNBC_T10,T16: raw data obtained from Navin *et al.* (2), processed by SCYN (3) and scSVAS offline pipeline. 10x_COLO829: raw data obtained from Velazquez *et al.* (4) and processed by scSVAS offline pipeline. |
| Embedding Map | TNBC_T10,T16 (2, 3), 10x_COLO829 (4) | TNBC_T10,T16: raw data obtained from Navin *et al.* (2), processed by SCYN (3) and scSVAS offline pipeline. 10x_COLO829: raw data obtained from Velazquez *et al.* (4) and processed by scSVAS offline pipeline. |
| Time Lineage | AML (5, 6), HGSOC_P7 (6, 7) | AML: raw data from Ding *et al.* (5), input files obtained from Smith *et al.* (6); HGSOCP7: raw data from McPherson *et al.* (7), input files obtained from Smith *et al.* (6) |
| Space Lineage | HGSOC_P1,P7 (6, 7), PC_A21 (6, 8) | HGSOC_P1, P7: raw data from McPherson *et al.* (7), input files obtained from Smith *et al.* (6)<br>PC_A21: raw data from Gundem *et al.* (8), input files obtained from Smith *et al.* (6) |
| Space Prevalence | HGSOC_P1,P7 (6, 7), PC_A21 (6, 8) | HGSOC_P1, P7: raw data from McPherson *et al.* (7), input files obtained from Smith *et al.* (6)<br>PC_A21: raw data from Gundem *et al.* (8), input files obtained from Smith *et al.* (6) |
| Clonal Lineage | TNBC_T10,T16 (2, 3), 10x_COLO829 (4) | TNBC_T10,T16: raw data obtained from Navin *et al.* (2), processed by SCYN (3) and scSVAS offline pipeline. 10x_COLO829: raw data obtained from Velazquez *et al.* (4) and processed by scSVAS offline pipeline. |
| Recurrent Event | LC | The lung cancer data present in Recurrent Event is prepared in another manuscript. |

## Supplementary Note 4: Supplementary Figures

**Fig. S1.**
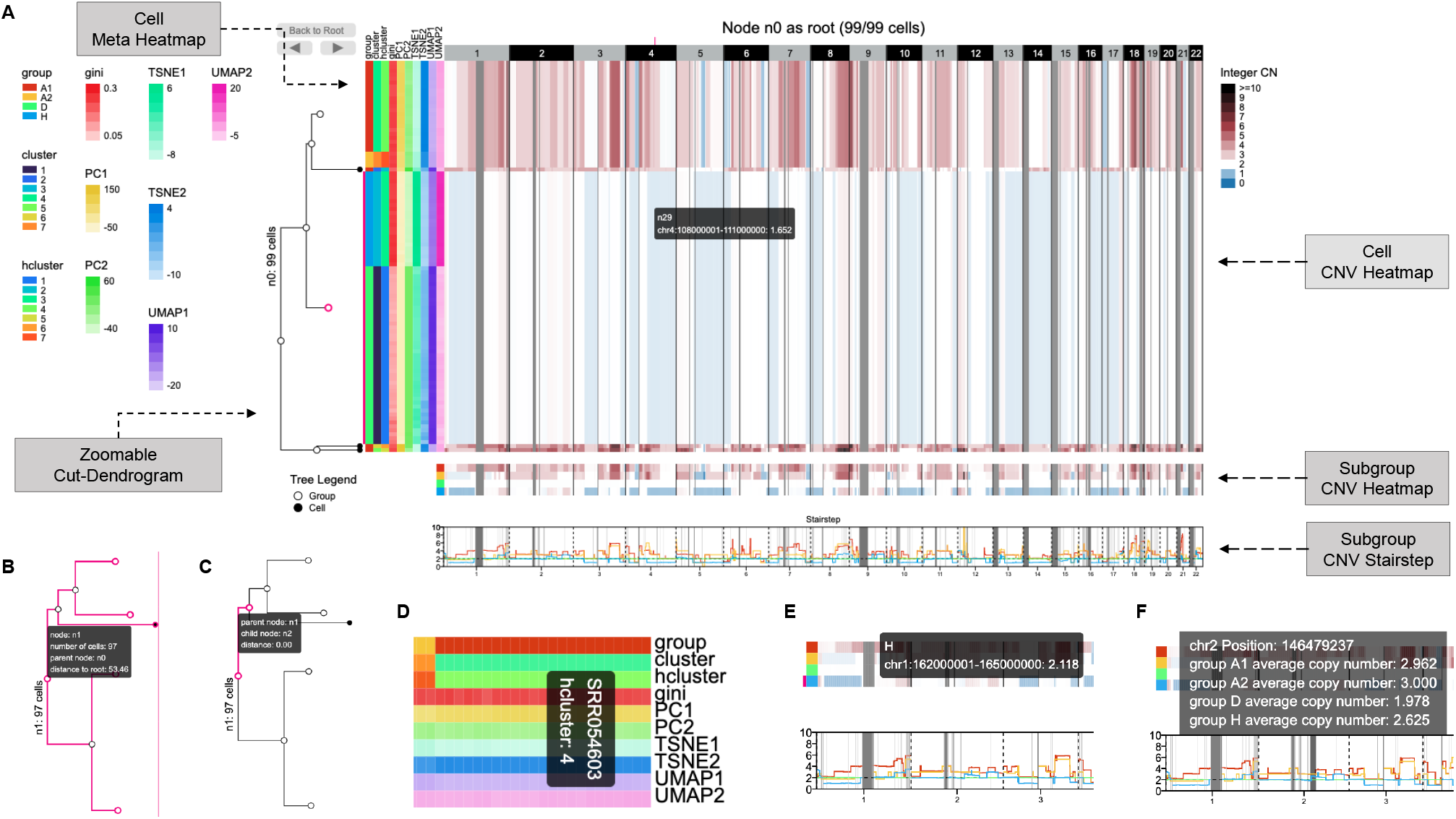
Demo representation of the CNV View interface. **A** The full display of CNV View. **B** The tooltip of zoomable cut dendrogram node. **C** The tooltip of zoomable cut dendrogram branch. **D** The tooltip of cell meta heatmap. **E** The tooltip of subgroup CNV heatmap. **F** The tooltip of subgroup CNV stairstep.

**Fig. S2.**
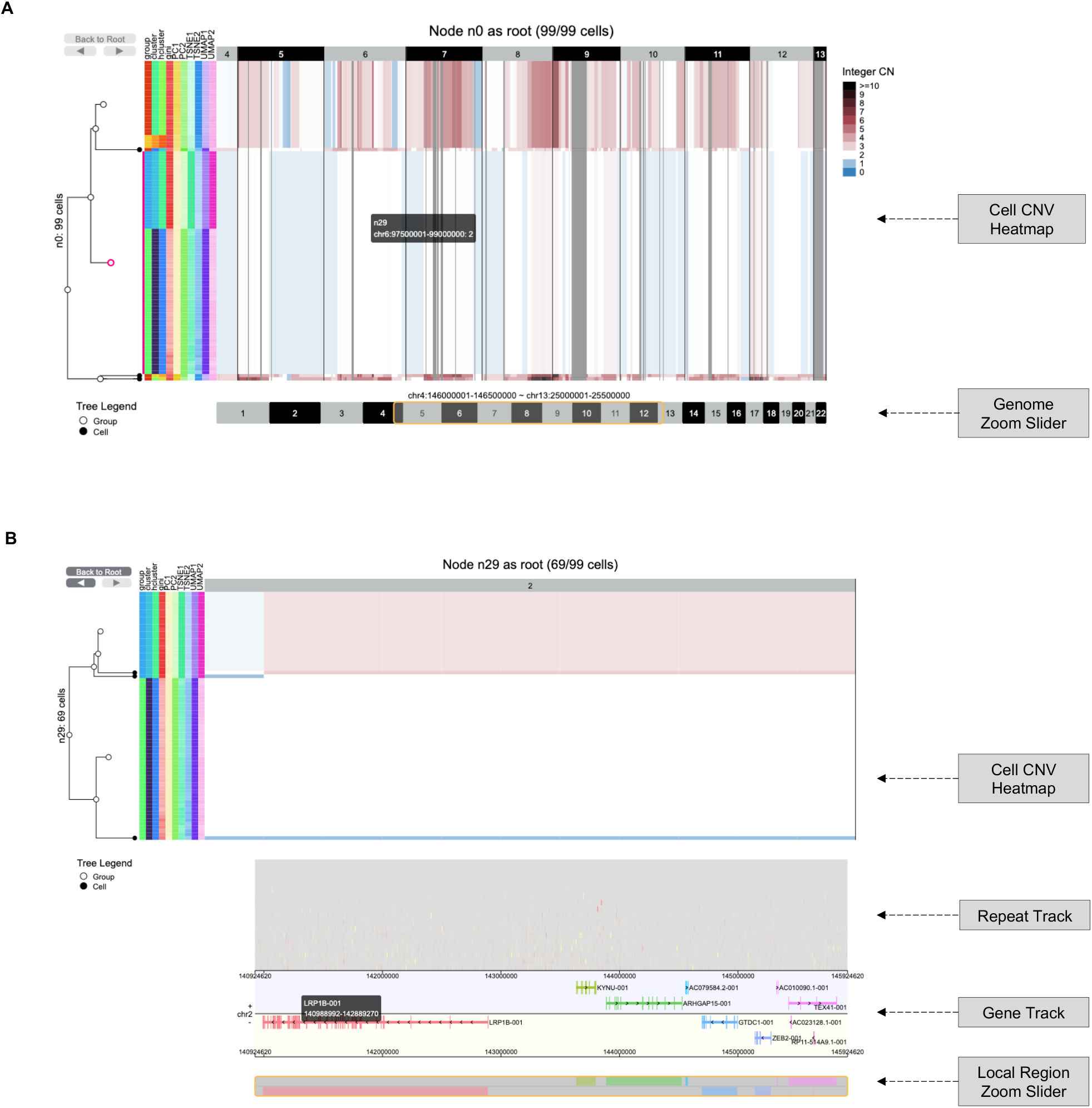
Demo representation of the CNV Heatmap interface. **A** The display of CNV Heatmap with genome zoom slider. **B** The display of local region CNV Heatmap with repeat and gene track.

**Fig. S3.**
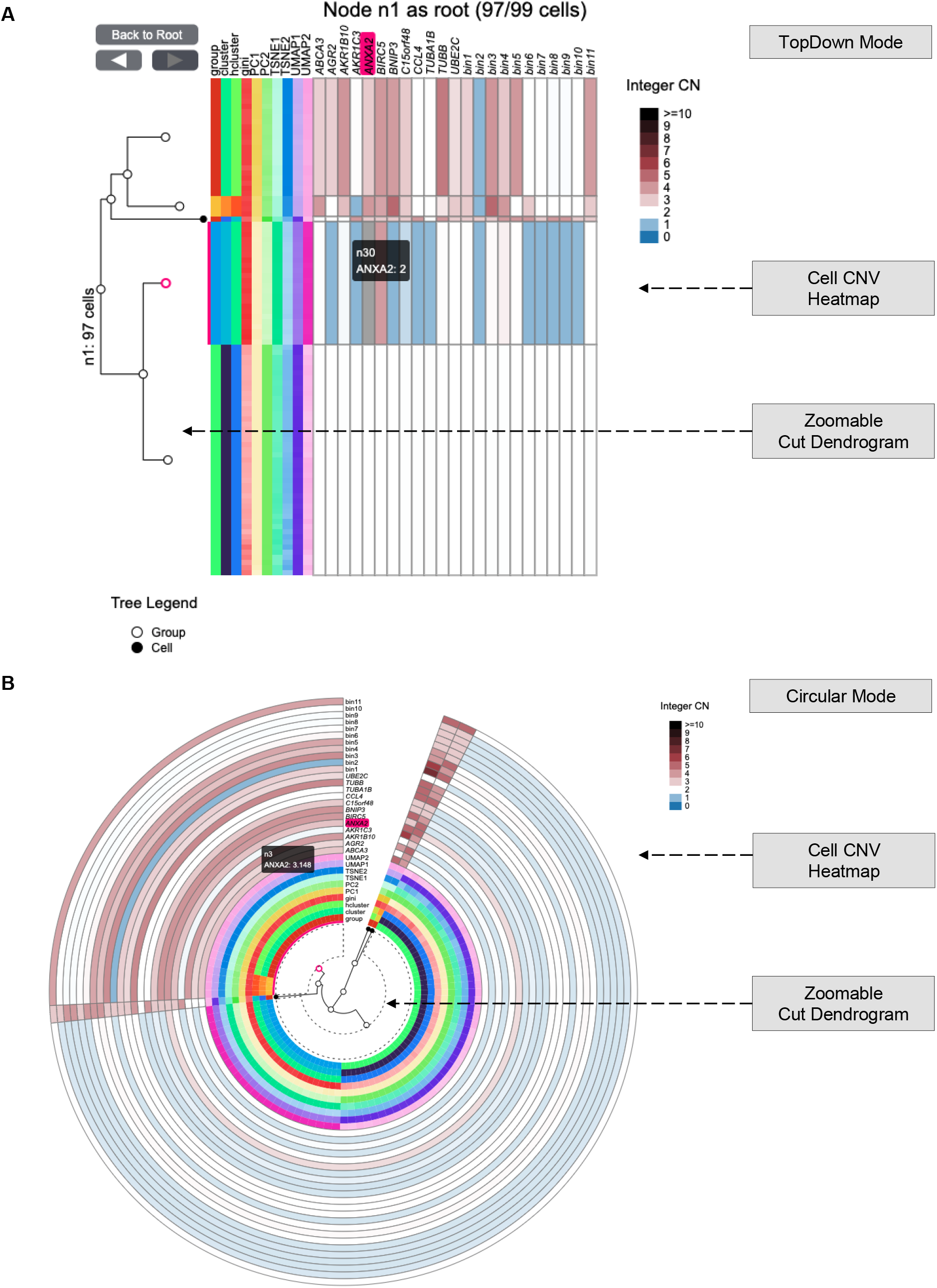
Demo representation of the Cell Phylogeny interface. **A** The display of Cell Phylogeny in topdown mode. **B** The display of Cell Phylogeney in circular mode.

**Fig. S4.**
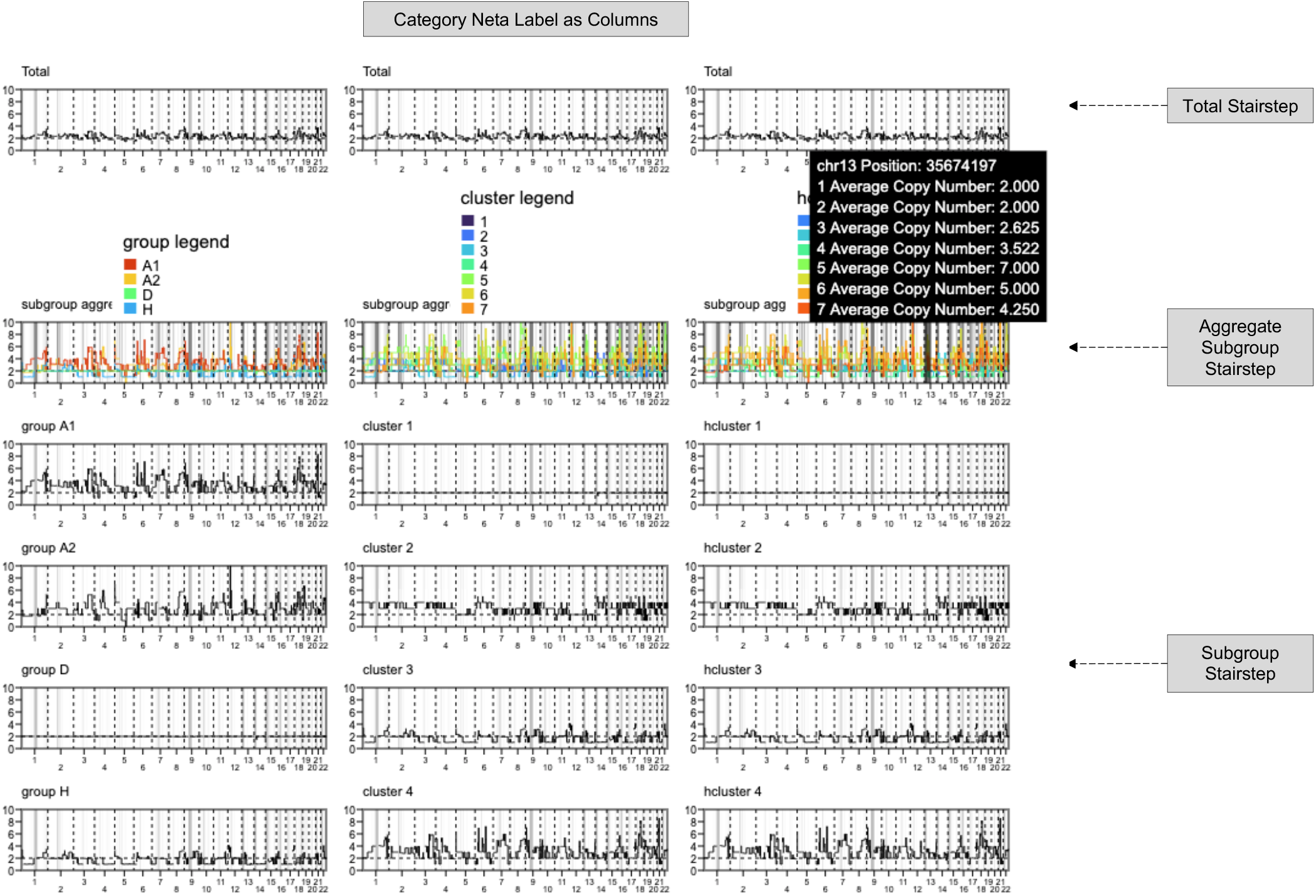
Demo representation of the Ploidy Stairstep interface.

**Fig. S5.**
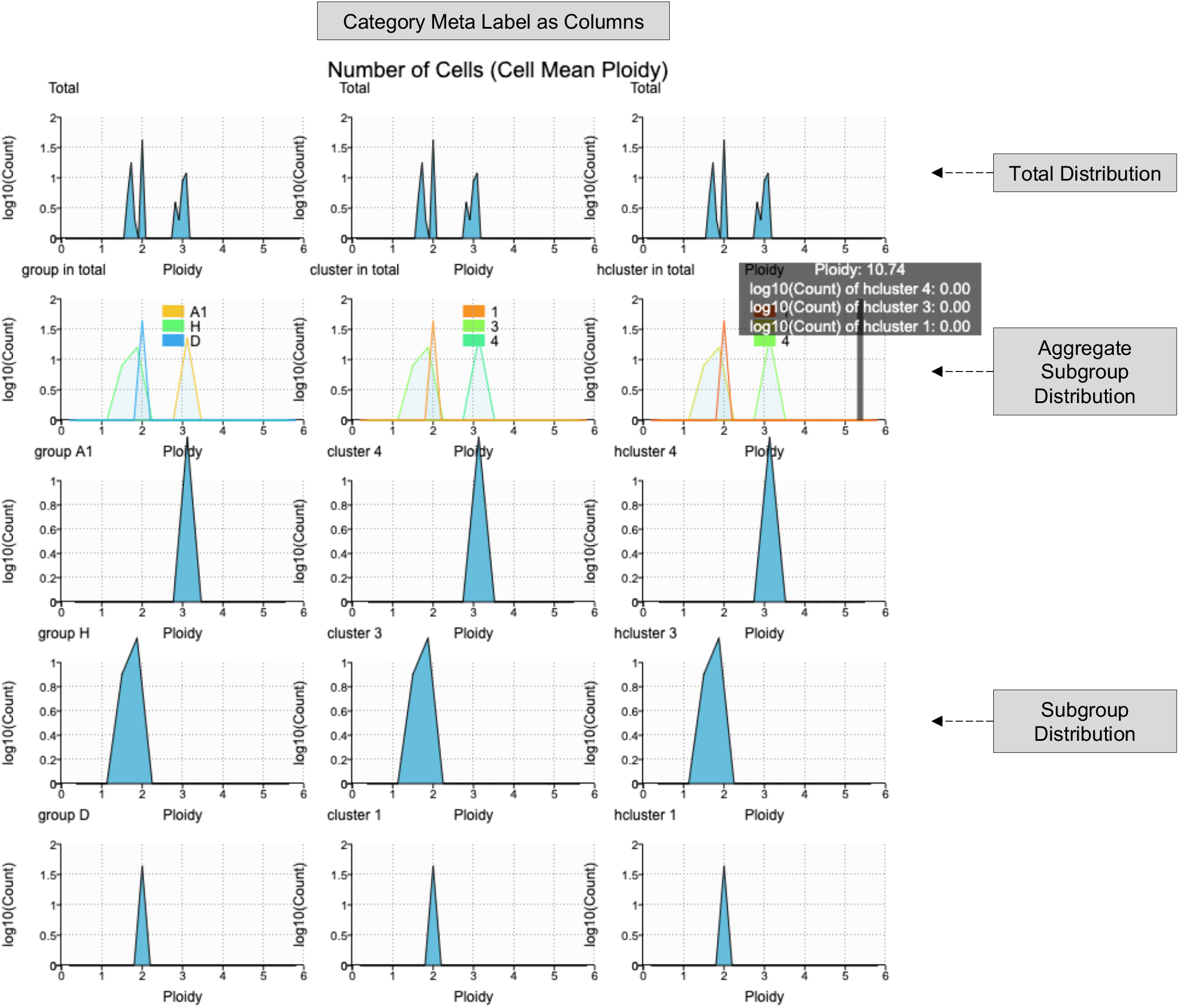
Demo representation of the Ploidy Distribution interface.

**Fig. S6.**
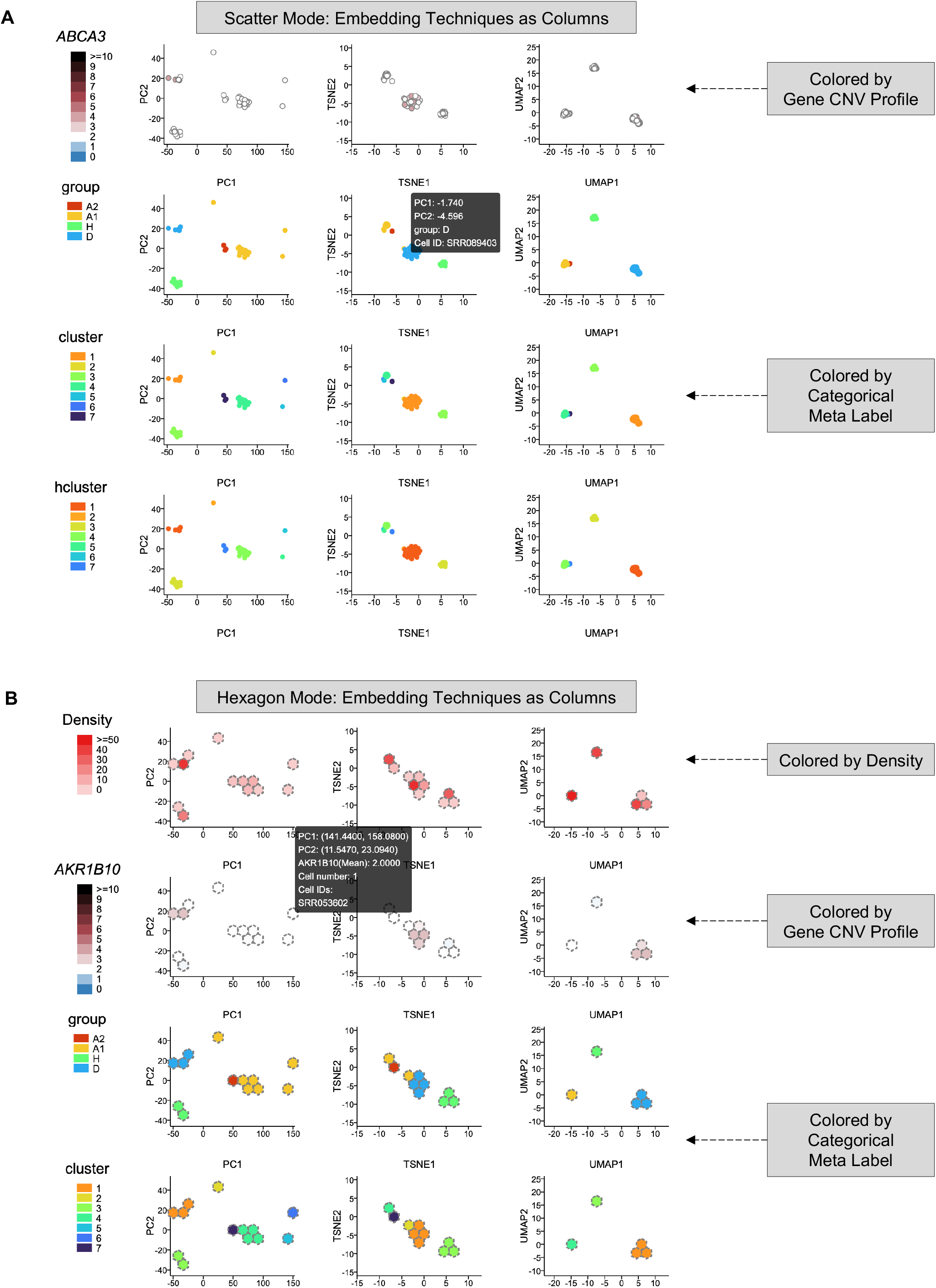
Demo representation of the Embedding Map interface. **A** The display of Embedding Map in scatter mode. **B** The display of Embedding Map in hexagon mode.

**Fig. S7.**
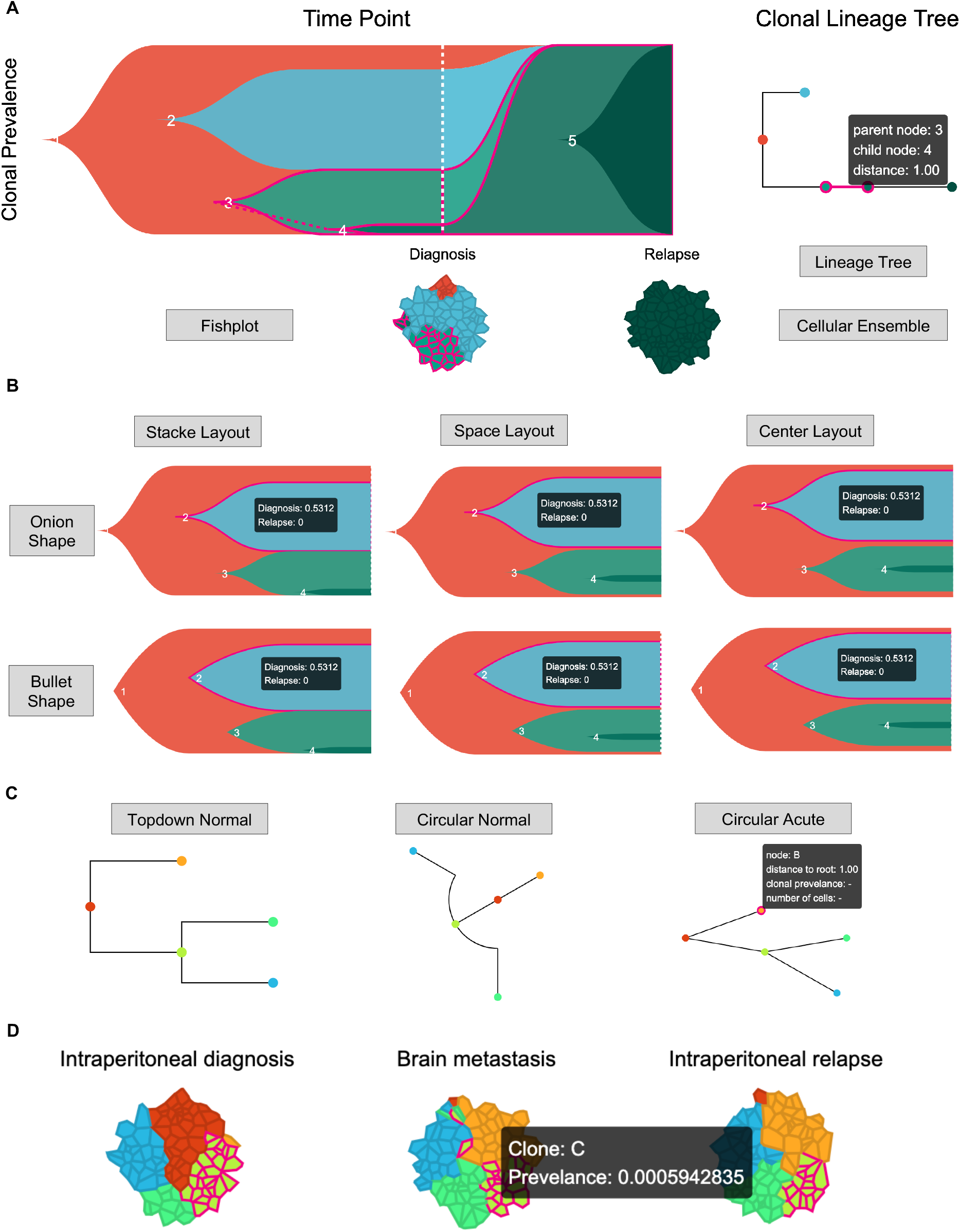
Demo representation of the Time Lineage interface. **A** The full display of Time Lineage. **B** The display of fishplot with available shape and layout. **C** The display of lineage tree with available shape. **D** The display of cellular ensemble.

**Fig. S8.**
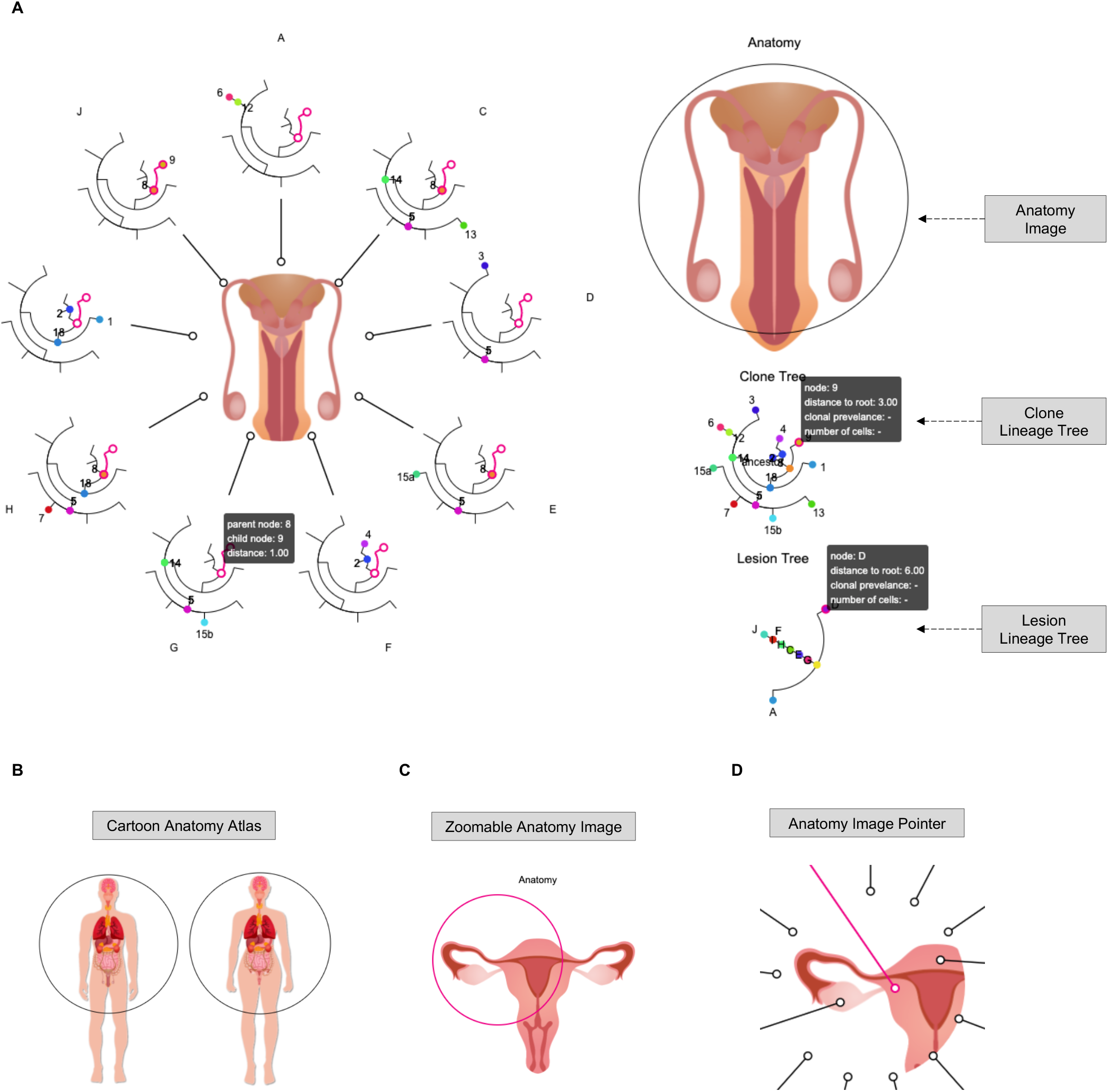
Demo representation of the Space Lineage interface. **A** The full display of Space Lineage. **B** Demonstration of cartoon anatomy atlas. **C** The display of zoomable anatomy image. **D** The display of anatomy image pointer. Freepik and macrovector / Freepik design the cartoon anatomy images, we acknowledge for the free license.

**Fig. S9.**
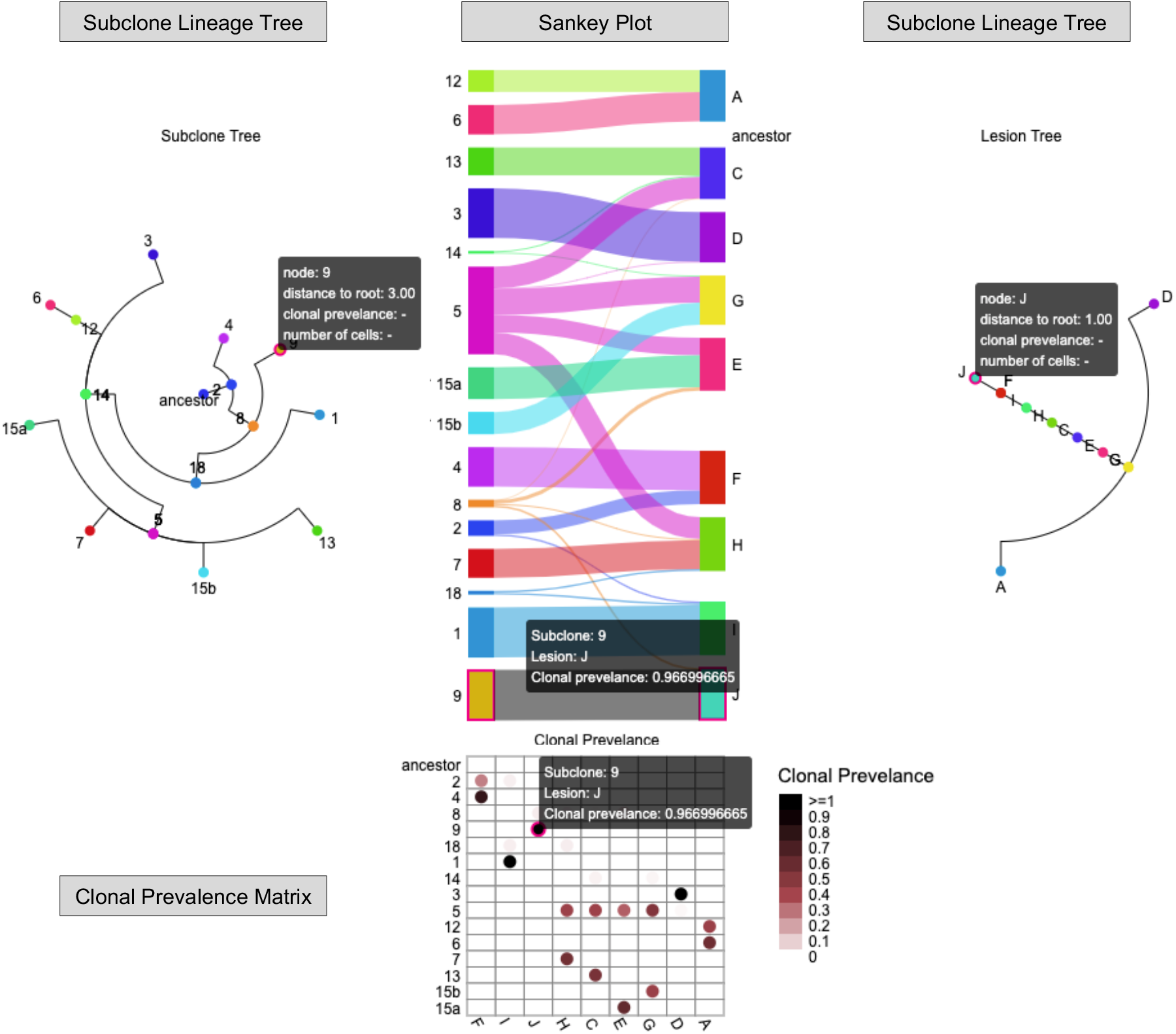
Demo representation of the Space Prevalence interface.

**Fig. S10.**
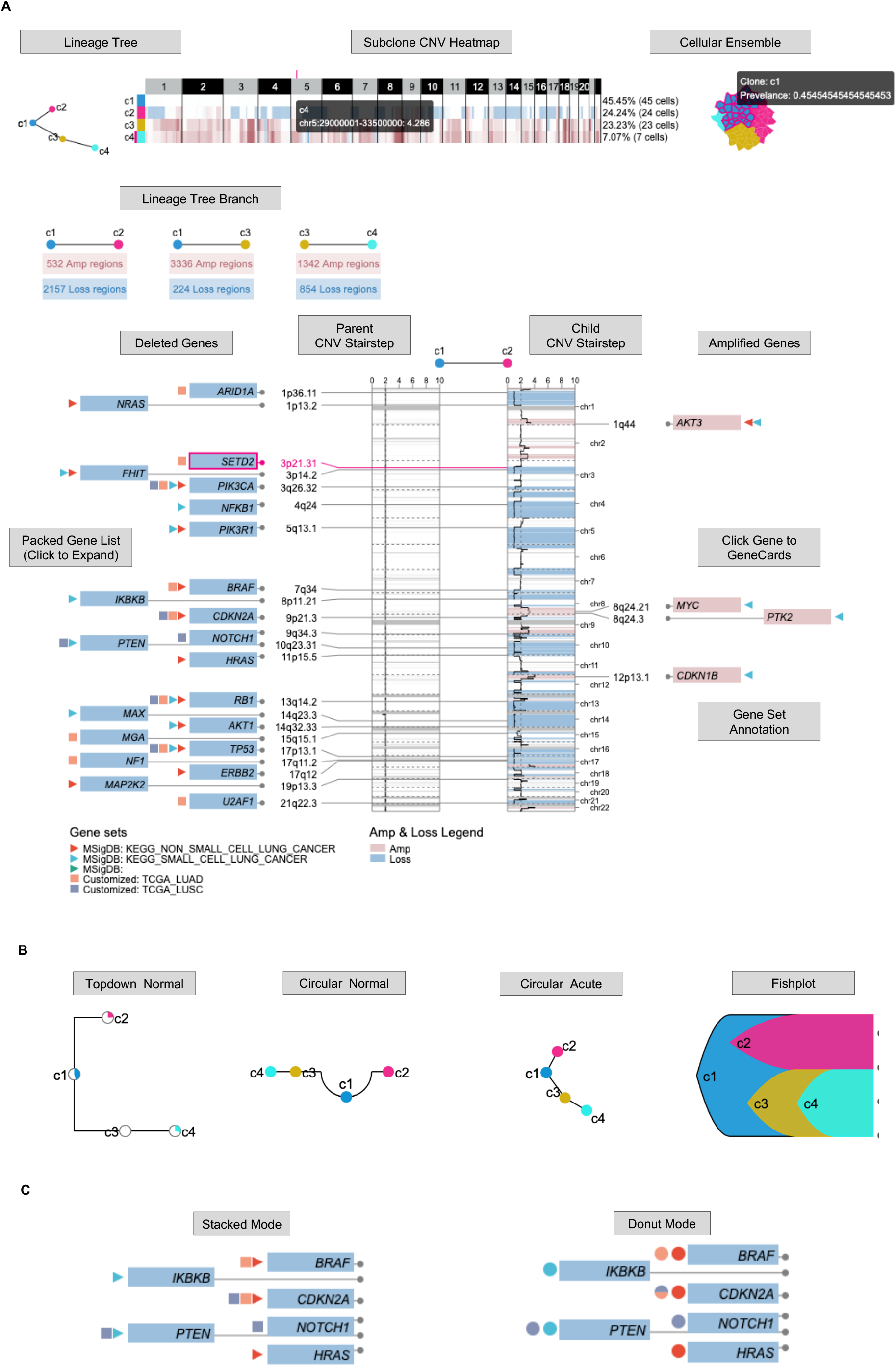
Demo representation of the Clonal Lineage interface. **A** The full display of Clonal Lineage. **B** The display of lineage tree with available shape. **C** The display of gene set annotation with stacked and donut modes.

**Fig. S11.**
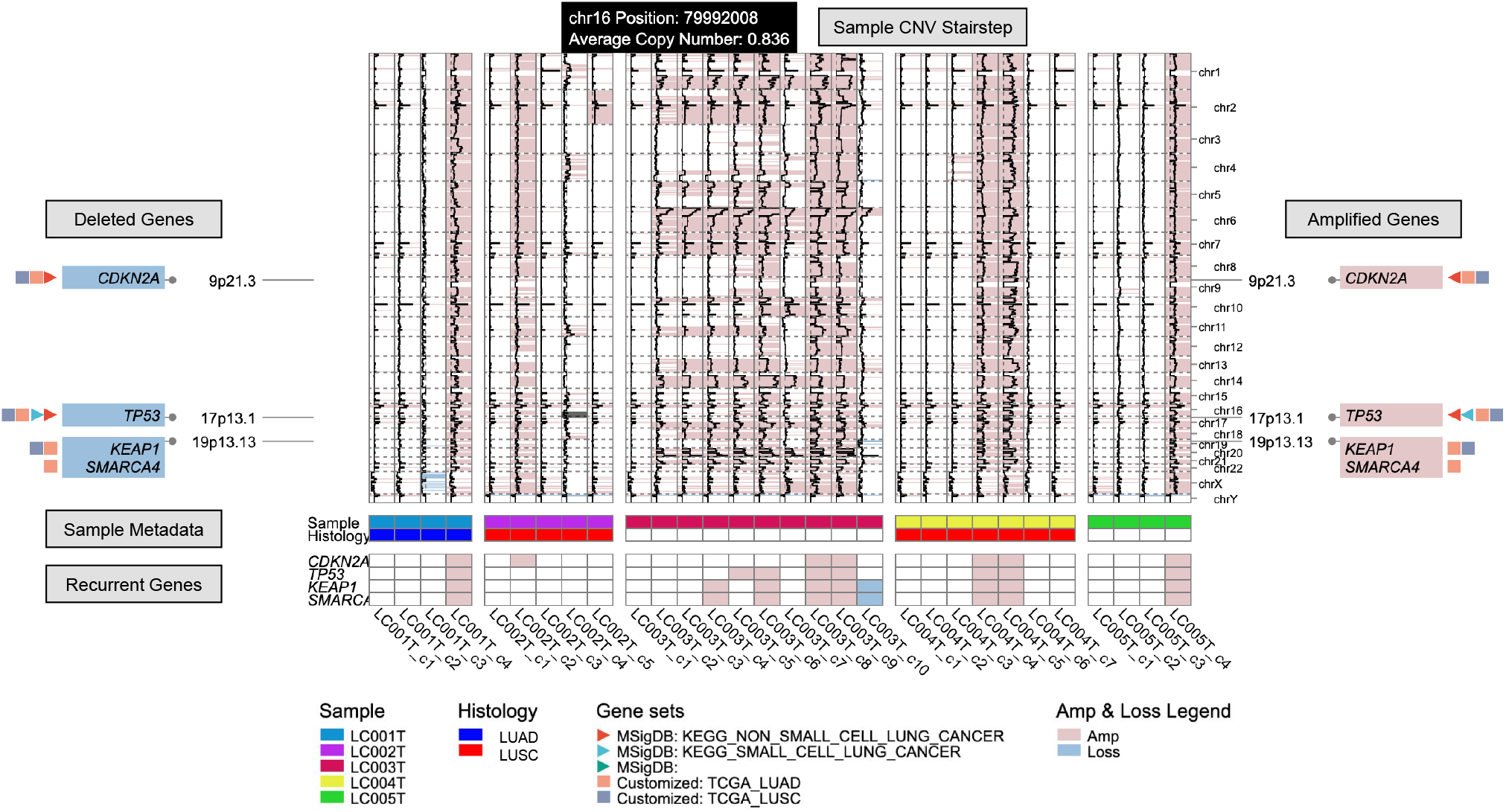
Demo representation of the Recurrent Event interface.

**Fig. S12.**
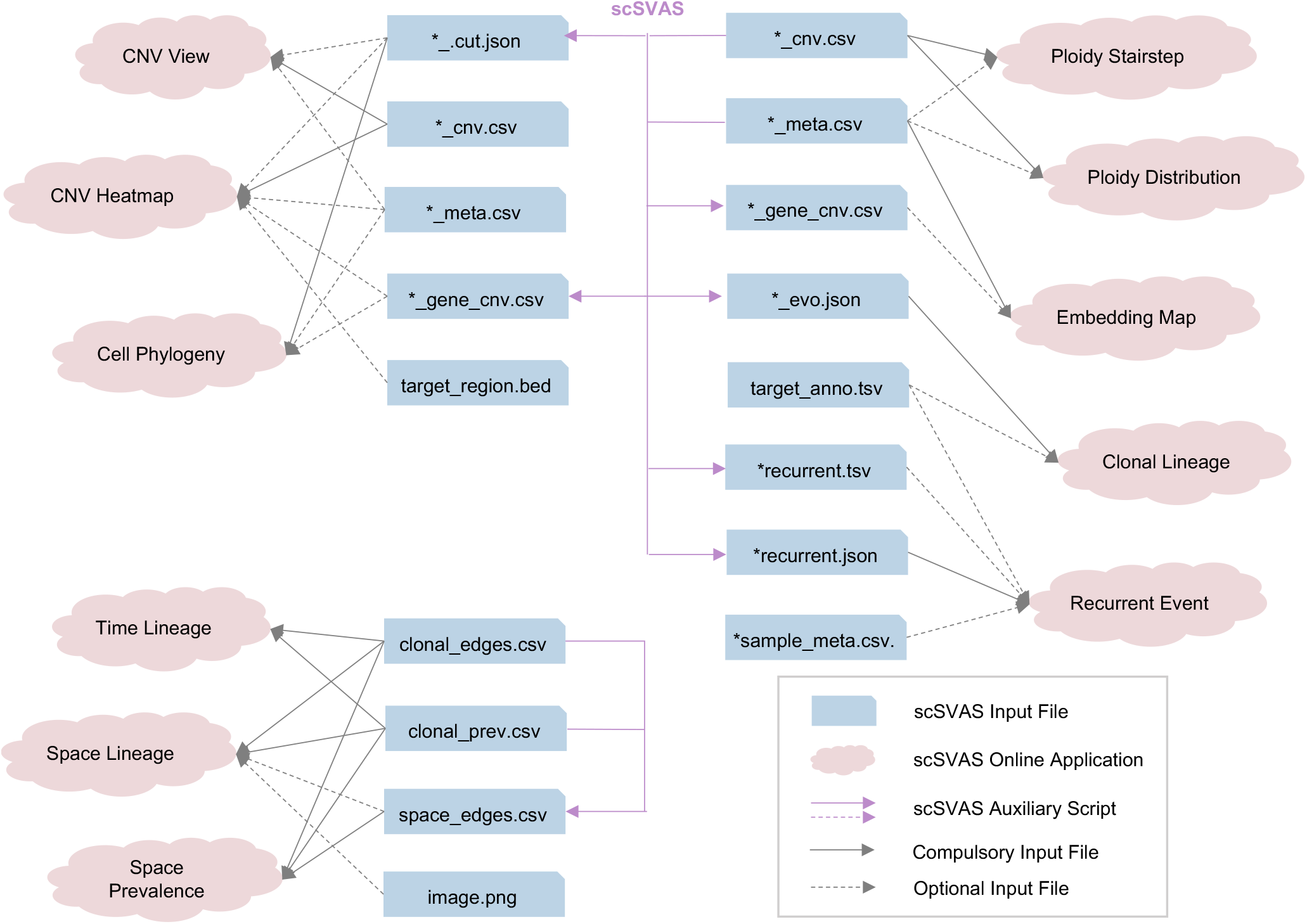
The input files required for scSVAS applications. The demo files are available at Editor in each application page and described in Table S2. The input format and offline pipeline are available at https://docsc.deepomics.org and https://github.com/paprikachan/scSVAS.

